# Immune cells adapt to distinct stem cell niches to govern tissue homeostasis

**DOI:** 10.64898/2026.01.28.701831

**Authors:** S. Martina Parigi, Sairaj M. Sajjath, Charlotte J. Bell, Shaopeng Yuan, Vitoria M. Olyntho, Alain R. Bonny, Sandra Nakandakari-Higa, Sergio A. Lira, Daniel Mucida, Gabriel D. Victora, Elaine Fuchs

**Affiliations:** Howard Hughes Medical Institute, Robin Chemers Neustein Laboratory of Mammalian Cell Biology and Development, The Rockefeller University, New York, NY USA; Howard Hughes Medical Institute, Laboratory of Mucosal Immunology, The Rockefeller University, New York, NY, USA; Precision Immunology Institute, Icahn School of Medicine at Mount Sinai, New York, NY 10029, USA; Howard Hughes Medical Institute, Laboratory of Lymphocyte Dynamics, The Rockefeller University, New York, NY, USA

## Abstract

In adult tissues, epithelial stem cells exist within distinct residences, each endowing them with exclusive instructions for regenerative fitness under homeostasis and stress. Key components of these ‘niches’ are immune cells, which classically protect the host against external and internal threats. Whether and how stem cell:immune cell crosstalk contributes to normal tissue biology remains less clear. Here, we discover functional adaptation of resident lymphocytes within two distinct skin stem cell niches and show that through this communication, each niche adjusts to meet diverse tissue demands. In the upper hair follicle, where microbial load is high, T cells express lymphotoxin-β and stimulate adjacent receptor-positive epithelial stem cells to form an immune-competent niche that controls microbial expansion. By contrast, in the epidermis, these T cells produce amphiregulin to maintain continuous stem cell reconstitution of the skin’s barrier. Concomitantly, they express the immune checkpoint protein ‘LAG-3’, which autorestricts lymphocyte numbers, and hence amphiregulin levels, thereby preventing over-proliferative responses. Finally, when epidermal T cells are absent, dermal lymphocytes restore the imbalance by colonizing and adapting to their new niche. Our findings unveil functional specialization and homeostatic resilience of immune-stem cell niches, each tailored to suit the demands of distinct tissue microenvironments.

## Main

Tissues are complex multicellular structures where the concerted action and interaction among different cell types, strategically positioned within spatially distinct microniches, ensures critical maintenance of a dynamic equilibrium called homeostasis^1,2^. Unveiling programs of cellular co-adaptation within geographically distinct cellular communities is thus essential to understand how tissue structure and function are preserved while being constantly rejuvenated. At the interface between the body and the outside world, the skin epithelium offers a unique model to study this question as it hosts a diverse array of cells organized in compartmentalized microniches within the tissue^3^.

At the core of this functional anatomic heterogeneity are the key regenerative units, tissue stem cells (SCs). In the adult, these SCs reside in discrete niches that through mechanisms still unfolding, endow them with their own tissue regenerative tasks and functions. In the skin, epithelial SCs in the innermost (basal) layer of the stratified interfollicular epidermis (IFE) undergo a continuous flux of lateral proliferation along the basement membrane at the epidermal-dermal border. This renewal is balanced by upward terminal differentiation to build and maintain a physical barrier at the skin surface that retains body fluids and protects the host from the external environment. Near the hair follicle (HF) orifices where hairs protrude from the skin surface, SCs also reside along this contiguous basement membrane, but here, their task is to mount a program that contains and adapts to bacterial colonization ^4–7^. Further down along the HF basement membrane are two epithelial SC compartments responsible for maintaining sebaceous glands and for driving hair regeneration during the hair cycle.

To sustain such diverse homeostatic functions, tissue SCs rely on the regional support of other cell types that are strategically positioned in close proximity (i.e. the stem cell niche)^3,8,9^. Tissue resident immune cells are prominent constituents of these niches, where they are classically tasked with protecting the host in the face of injuries and other environmental threats^10–14^. Whether immune cells acquire homeostatic programs when colonizing a SC niche, and as importantly, whether these programs become tailored to the specialized needs of their immediate geography, remain largely unknown. Here, we show that skin SCs and immune cells co-adapt within distinct tissue microniches and that disruption of their crosstalk geographically perturbs tissue homeostasis. Additionally, we unearth adaptive strategies that have evolved to maintain key SC-immune cell signalling within homeostatic niches and to employ compensatory mechanisms to reinstate them when perturbed and thereby maintain the equilibrium of the tissue.

### Immune cell transcriptomes are influenced by direct contact with epithelial SCs residing in distinct niches

To interrogate whether immune cells are imprinted with specific functional features within SC niches, we leveraged the recently developed technology, namely “Universal Labelling Immune Partnership by SorTagging Intercellular Contacts’, or uLIPSTIC^15^. This mouse model exploits the ability of *S. aureus* transpeptidase sortase A (SrtA) to covalently transfer a biotin-labelled substrate with sorting motif “LPTXTG” to a physically juxtaposed N-terminal oligoglycin (G5) (**Fig. 1a**). In uLIPSTIC mice, the recipient machinery (i.e. G5) is ubiquitously expressed, while the donor machinery (i.e. SrtA) is preceded by a floxed STOP codon, excised only in CRE-recombinase expressing cells (**Extended Data Fig. 1a**).

**Figure 1.**
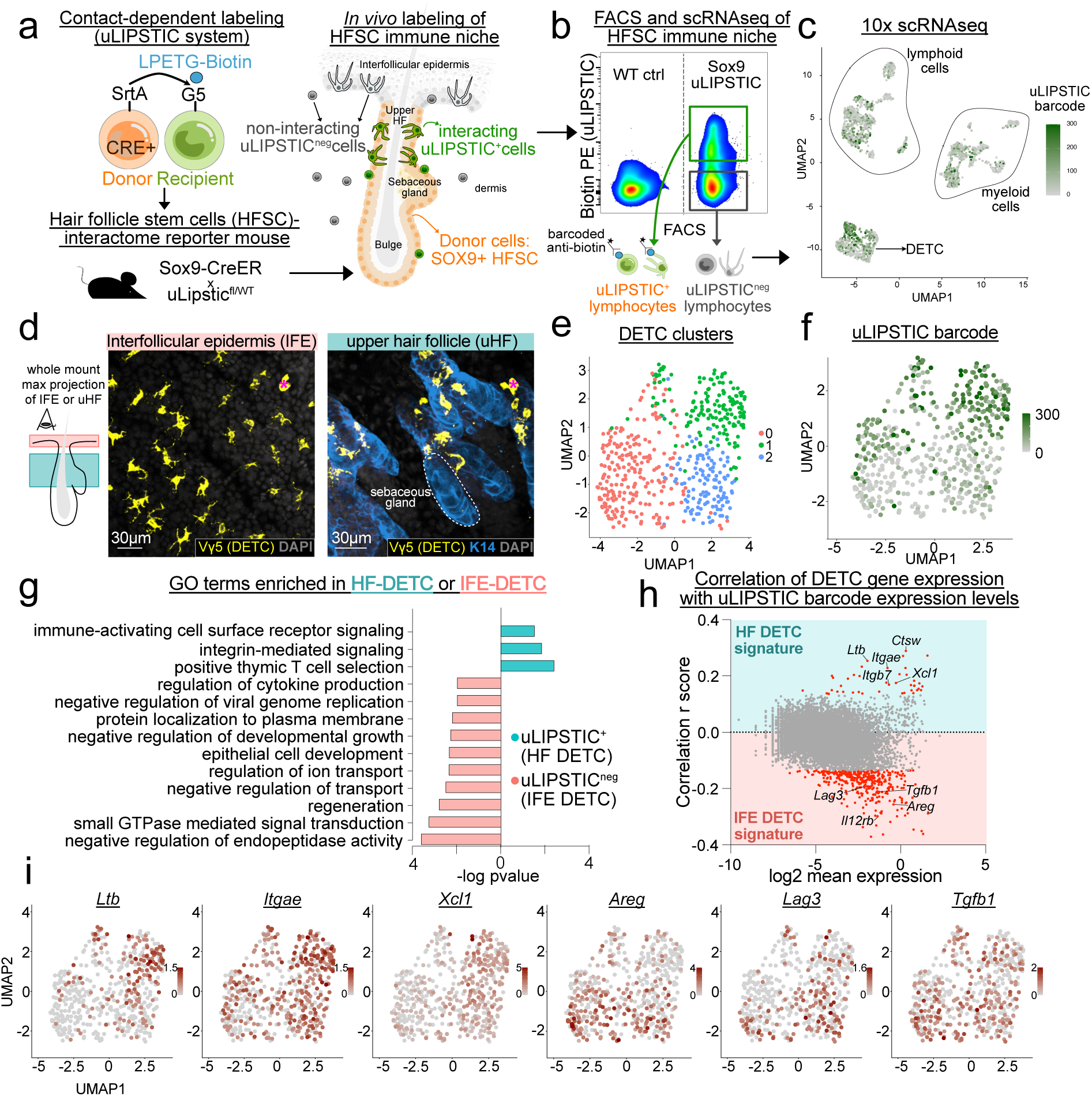
Unbiased transcriptional profiling of immune:SC niche interactions within the skin epithelium. **a**, uLIPSTIC system designed to transfer biotin to recipient cells that directly interact with *Sox9-CreER* expressing hair follicle stem cells residing within discrete niches in the upper hair follicle (uHF), sebaceous gland and bulge. **b,** Focusing on immune:SC interactions, representative FACS plot of biotinylated versus unlabelled immune cells in Sox9-uLIPSTIC compared to WT skin. These populations were single-cell sequenced and HF SC-interacting cells were distinguished by barcoded anti-biotin antibodies. **c,** Uniform Manifold Projection (UMAP) plot showing major immune cell types labelled in green by abundance of uLIPSTIC barcode reads. **d,** Whole mount Vγ5 immunofluorescence images in a max-projection, depicting DETCs prominence in the interfollicular epidermis (IFE) and in the uHF. Scale bar 30μm. * denote autofluorescence. **(e-f)** UMAP plots color-coded based on unbiased Seurat clustering analysis of DETCs **(e)** and uLIPSTIC barcode reads **(f)**. **g,** barplot of GeneOntology biological processes enriched in uLIPSTIC^+^ (HF-DETC) or uLIPSTIC^neg^ (IFE-DETC). **h,** volcano plot showing intensity of gene expression (log2 mean gene expression) and correlation score of each gene with uLIPSTIC barcode reads. Red dots denote transcripts whose differential expression is statistically significant (false discovery rate <0.05). **i,** Representative transcripts enriched in IFE- or HF-DETC were displayed on DETC UMAP plots, color-coded based expression levels. Five *Sox9-uLIPSTIC* mice in second telogen were pooled into a single sample for sequencing. Further details on statistics and reproducibility in Methods. See Extended Data Fig. 1 for additional supporting experiments.

To distinguish immune cells within the SC niches of the hair follicle from those within the IFE, we generated uLIPSTIC x *Sox9-CreER* mice (i.e. hair follicle SC interactome reporter mice, **Fig. 1a**)^16^. During the resting phase (telogen) of the hair cycle, the donor machinery was selectively expressed by SOX9^+^ SCs residing in the niches of the upper hair follicle (uHF), sebaceous gland and bulge (the SC niche governing the hair cycle) (**Extended Data Fig. 1b-c**). The biotin-LPETG substrate was transferred to uLIPSTIC^+^ cells in direct physical contact with the SOX9-expressing SCs (**Fig. 1a**). With this strategy, the CD45^+^ immune cells in the HF niches became biotin-labelled, while those within the IFE or dermis remained unlabelled.

To isolate and characterize the biotin-labelled immune interactor(s), we stained with anti-biotin barcoded antibody, sorted by flow-cytometry activated cell sorting (FACS) and processed for 10X single-cell RNA sequencing. We additionally captured the biotin^neg^ immune cells, which encompassed immune cells in contact with SOX9^neg^ IFE-SCs and non-interacting immune cells (like the ones found in the dermis) (**Fig. 1b-c**). To ensure proper representation of diverse niche components, we sequenced comparable numbers of biotin^+^ and biotin^neg^ lymphoid and myeloid immune cells (**Extended Data Fig. 1d-e**).

Skin epitheliotropic T cells are often located around the upper hair follicle, where they express Integrin αE, enabling them to form heterophilic cell adhesions with E-cadherin on the surface of the skin epithelium^15,17^. Indeed, we observed a robust positive correlation between uLIPSTIC barcode (biotin^+^) signal and *Itgae* expression in both CD4^+^ and CD8^+^ T lymphocytes (**Extended Data Fig. 1f**). Similarly, uLIPSTIC-barcoded innate lymphoid cells (ILCs) expressed several genes (e.g. *Capg, Ccr6* and *Ctsw*) that had previously been assigned to ILCs^18^ around the sebaceous glands. These results provided compelling evidence that our uLIPSTIC strategy efficiently and selectively labelled immune cells within hair follicle SC niches.

Realizing that the uLIPSTIC approach could provide us with the unique opportunity to interrogate whether the same immune cell tailors its behaviour to suit the particular needs of its SC niche, we focused on dendritic epidermal T cells (DETCs). This subpopulation of lymphocytes expresses high levels of the invariant Vγ5Vδ1 T cell receptor and participates in wound healing and tumour immune-surveillance^19–25^. As exclusively intraepithelial immune cells, any DETCs that did not label by interacting with SOX9^+^ HF-SCs were thus likely to reside in the IFE-SC niche, thus representing an ideal cellular prototype to address whether virtually identical cells are able to differentially adapt within distinct micro-niches.

Whole mount immunofluorescence with antibodies against Vγ5 further revealed that within the HF, DETCs were concentrated in the uHF, whose lower boundary was just above the sebaceous gland (**Fig. 1d**). At this location and within the IFE, DETCs intercalated among the stem cells. As shown by FACS analysis, the DETCs in both uLIPSTIC^+^ and uLIPSTIC ^neg^ fractions were typified by their high TCRγδ expression and their specific expression of Vγ5^+^ (**Extended Data Fig. 1g**).

DETCs were transcriptionally heterogeneous, in agreement with a previous report^26^ (**Fig. 1e**). Strikingly, however, by overlaying these cells’ gene expression profiles with their uLIPSTIC barcode signal, we discovered that this heterogeneity is rooted in their distinct geographical positionings in the skin. Thus, the uLIPSTIC^+^ DETCs within the upper HF SC niche (uHF-DETCs) displayed a transcriptome that was distinct from that of uLIPSTIC^neg^ DETCs within the IFE SC niche (IFE-DETCs) (**Fig. 1f**).

uHF-DETCs were enriched for pathways such as ‘immune-response activating receptor signalling’ and ‘integrin-mediated signalling’, while IFE-DETCs featured processes such as ‘regulation of epithelial cell development and regeneration’ (**Fig. 1g**). Delving deeper into specific genes mediating these different programs, we binned DETC genes according to those that either positively or negatively correlated with their uLIPSTIC barcode signal. Intriguingly, uHF-DETCs showed pronounced expression of *Ltb, Itgae, Xcl1* and *Ctsw*, which positively correlated with uLIPSTIC barcode intensity, while IFE-DETCs were enriched for *Areg, Lag3, Tg@1* and *il12rb,* which anti-correlated with uLIPSTIC barcode intensity (**Fig. 1h-i**). Altogether, these results showcased the power of spatial sequencing technologies when based upon direct cell-cell contact. Our data led us to posit that through direct interactions with different skin epithelial SCs, the same immune cell type (here the Vγ5Vδ1^+^ T lymphocyte) may be able to fine-tune its transcriptome.

### DETCs acquire specific programs within distinct SC niches

To begin to assess whether the niche-dependent transcriptional differences of DETCs represent a necessary adaptation to sustain the distinct functions of uHF- and IFE stem cells, we focused our analysis on genes encoding DETC ligands that might differentially stimulate immune crosstalk with receptors expressed by skin epithelial SCs^27–30^. Notable among these were *Ltb,* encoding lymphotoxin-β (LTB); *Areg,* encoding Amphiregulin (AREG) and *Lag3,* encoding Lymphocyte Activation Gene 3 (LAG-3).

In skin at steady state, *Ltb* transcripts were enriched in several lymphocyte subtypes, including the DETCs (**Fig. 2a** and **Extended Data Fig. 2a-b**). In the absence of a validated FACS antibody to detect LTB in uLIPSTIC mice, we crossed *R26-tdTomato* mice with mice expressing CRE recombinase under the endogenous promoter of *Ltb* (*Ltb-EGFP-F2A-CreER*) (**Fig. 2b**) and confirmed by immunofluorescence significant enrichment of LTB^+^ DETCs in the uHF compared to the IFE (**Fig. 2c**). Intriguingly, even the other LTB-expressing cells, including dermal T lymphocytes and ILCs, were predominantly restricted to the uHF region (**Fig. 2c** and **Extended Data Fig. 2c**).^18^

**Figure 2.**
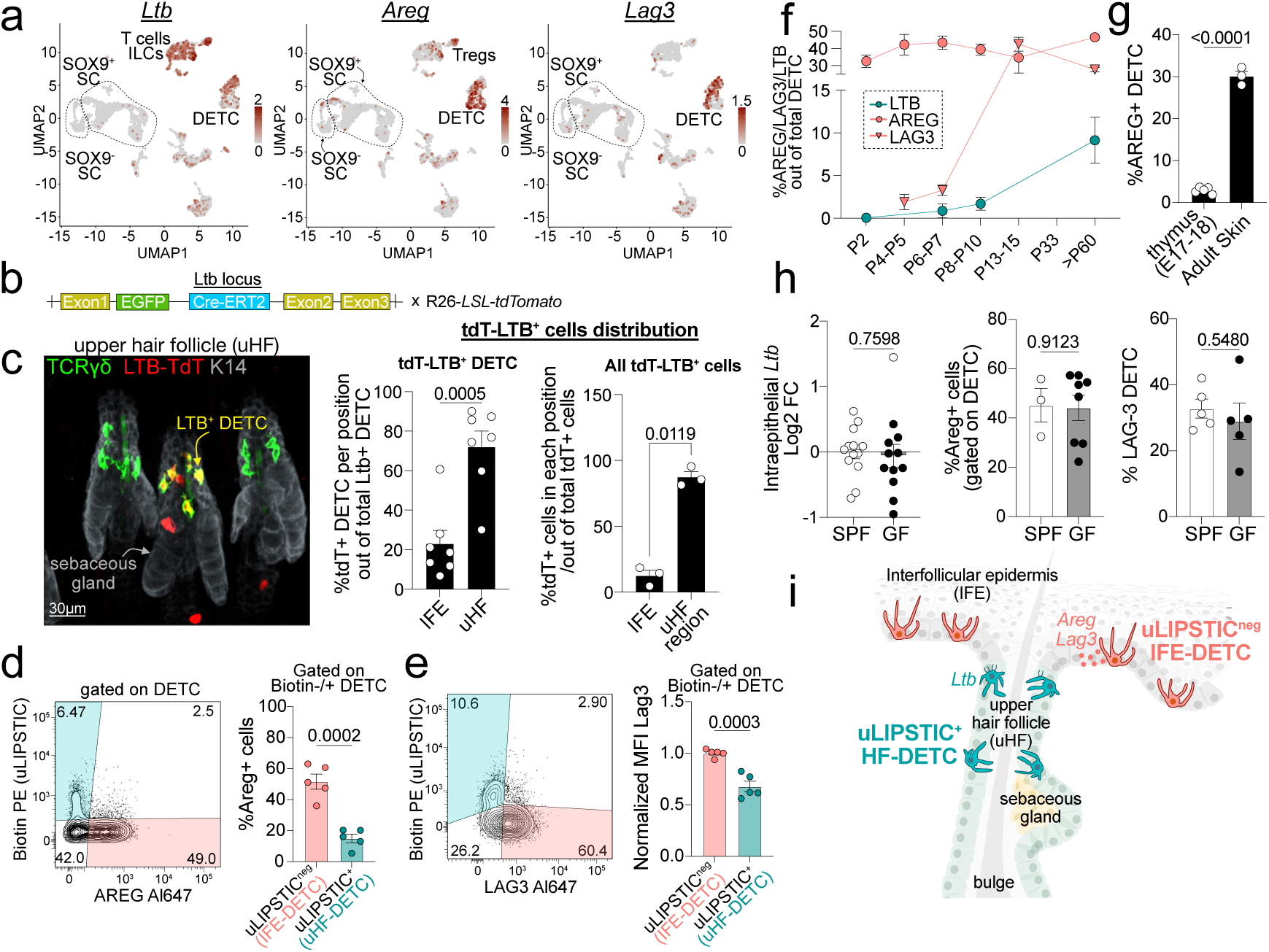
DETC expression of AREG, LAG-3 and LTB in the homeostatic skin is compartmentalized to distinct stem cell niches. **a**, UMAP plots displaying distribution of *Ltb, Areg* and *Lag3* transcripts across different cell types in mouse skin. Each dot is one cell, color-coded based on transcript expression levels. **b,** Schematic of the *Ltb-CreER x R26-tdT* mouse line. **c,** Left: Representative whole mount immunofluorescence showing that double positive (tdT^+^ and TCRγδ^+^) LTB-expressing DETCs reside primarily within the uHF compartment as do other LTB-expressing lymphocytes (tdT^+^ and TCRγδ^neg^). Epithelial SCs are marked by K14. Scale bar 30μm. Right: quantifications of (left) the probability of finding LTB^+^ DETCs (tdT^+^ and TCRγδ^+^) in uHF vs IFE and (right) probability of finding any LTB^+^ cell (tdT^+^) in the uHF region vs IFE. Each circle represents data from one mouse (3-8 fields of view). **d-e**, Representative FACS plots and quantifications showing that most DETCs expressing AREG **(d)** and LAG-3 **(e)** are uLIPSTIC^neg^ (IFE-DETC) and not uLIPSTIC^+^ (uHF-DETC). MFI, mean fluorescence intensity. **f,** Percentage of LTB-tdT^+^, AREG^+^ or LAG-3^+^ DETC out of total DETC as quantified by FACS of the skins of mice at the indicated postnatal time points. Each point represents the mean with SEM of 2-7 mice. **g,** Percentage of AREG^+^ DETC in the embryonic thymus of E17-18 pups and from the back skin of adult second telogen mice as quantified by FACS. **h,** Quantifications of total skin intraepithelial *Ltb* transcripts determined by qPCR, and AREG^+^ or LAG-3^+^ DETC determined by FACS when mice are kept in specific pathogen free (SPF) or germ-free (GF) facility. **i,** Schematic summarizing the compartmentalization of the two DETC niche programs in steady state. Data are representative of two-four (panel h), two (panel e), three (panel c, d, g) or three-seven (panel f) independent experiments and, unless indicated, each circle represents one mouse. Data in c, d, g and h are analysed by unpaired two-tailed Student’s *t-*test. *p-*values are indicated in each figure and data are represented as mean with SEM. Further details on statistics and reproducibility in Methods. See Extended Data Fig. 2 for additional supporting experiments.

In contrast to *Ltb*, *Areg* and *Lag3* were mainly expressed by DETCs within the interfollicular epidermis (**Fig. 1h-i** and **2a**). FACS analysis of AREG and LAG-3 in the uLIPSTIC HF-SC reporter interactome mice confirmed that the levels of these two ligands were significantly higher in IFE-DETC (uLIPSTIC^neg^) than in the uHF-DETC (uLIPSTIC^+^) (**Fig. 2d-e**).

### Transcriptional reprogramming of DETCs takes place after their arrival to the skin

DETCs originate in the thymus around embryonic day (E)14-15 and begin seeding the skin around E16-18^31^. As the thymus matures, DETCs are positively selected by binding their cognate antigen SKINT1. During this developmental window, DETCs become pre-programmed to home to the skin epithelium, which also expresses this protein^26,32–36^.

Our results thus far suggested that the epithelial stem cells of the IFE and the uHF may weave distinct niches that further shape the transcriptomes of their immune cell residents, in this case, DETCs. If so, these programs should only emerge *after* the DETCs migrate from the thymus and take up residence within the skin. To test this hypothesis, we analysed the kinetics of LTB, AREG and LAG-3 appearance in DETCs. Interestingly, expression of LTB and LAG-3 in DETCs did not begin until around postnatal day (P) 5-10, i.e. long after their maturation in the thymus and their migration into the skin (**Fig. 2f**). AREG was expressed from P2 onward in skin-resident DETCs and remained constant throughout later homeostasis (**Fig. 2f**). However, even for AREG, thymic DETC precursors were largely absent for this ligand, suggesting that the skin microenvironment was driving these distinctive ligand signatures in DETCs (**Fig. 2g**).

Upon birth, mice are exposed to the external environment, including the skin’s microbiota that then takes residency within the first several days of life^37^. However, DETC abundance and distribution within the IFE and uHF regions were unaffected by commensal colonization (**Extended Data Fig. 2d-e**). This differed from dermal TCRγδ^dim^ RORγt^+^ T cells, which are known to be microbiota-dependent^38^, and whose IL-17 production was significantly lower in germ-free (GF) versus specific pathogen free (SPF) mice (**Extended Data Fig. 2f**). Although the lack of specific antibodies precluded analysing the effects of the microbiota on LTB protein, total levels of intraepithelial *Ltb* transcripts were not markedly impacted (**Fig. 2h**). Similarly, AREG and LAG-3 proteins were expressed comparably in DETCs irrespective of commensal colonization (**Fig. 2h and Extended Data Fig. 2g**). Overall, our results revealed that during the emergence of specialized epithelial SC niches within the developing skin, DETCs home to these niches, where they are reprogrammed. This reprogramming occurred concomitantly, but was not necessarily influenced by, microbial colonization (**Fig. 2i**). Whether such programs are tailored to the SCs within these different niches and whether they are required to promote their respective homeostasis remained to be explored.

### Immune:SC specific LTB-LTBR signalling drives an immune-competent niche and restrains bacterial colonization at the HF orifice

Our studies thus far revealed that under homeostatic conditions, the uHF microniche is enriched with lymphocytes that express *Ltb* (**Fig. 2a**). Moreover, although *Ltbr* encoding its cognate lymphotoxinβ receptor (LTBR)^28^, was expressed more broadly, its expression and that of its downstream effector, *Nffib2,* were markedly pronounced in the uHF-SCs (**Extended Data Fig. 3a-b**). These data suggested a special importance of LTB-LTBR mediated immune:SC signalling within the uHF SC compartment.

To address the downstream consequences of LTB signalling within the uHF microenvironment, we treated mice intradermally with an LTBR agonist or an isotype as control. Following FACS purification of the uHF-SCs, we performed RNA sequencing (**Fig. 3a** and **Extended Data Fig. 3c**). LTBR-stimulated uHF-SCs robustly induced transcripts involved in inflammatory responses, T cell activation and response to bacteria (**Extended Data Fig. 4a**). Among these genes were lymphocyte-recruiting chemokines, including *Cxcl10* and *Cxcl16* (**Fig. 3a**).

**Figure 3.**
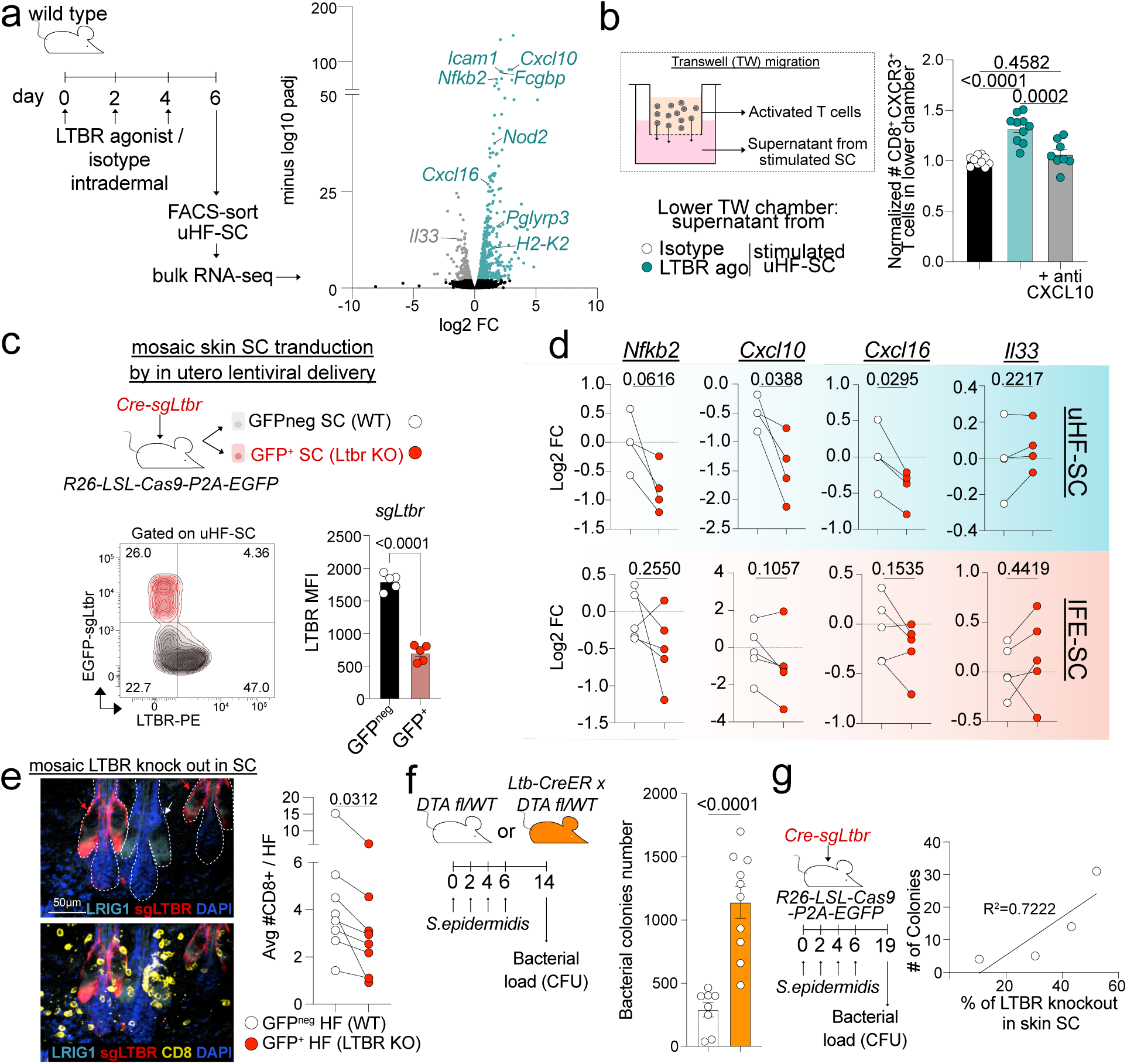
Immune: stem cell interactions within the uHF niche drive LTB-LTBR signalling and downstream lymphocyte recruitment and protection against bacterial colonization. **a**, Left: strategy to elevate LTBR signalling in uHF-SCs. Right: volcano plot showing examples of genes up- and down-regulated in uHF-SC upon intradermal administration of LTBR-agonist (compared to isotype control) in mice. Two mice/condition were sequenced. **b,** Left: strategy of the *in vitro* transwell experiment to test CD8^+^ T cell migration towards the supernatant of uHF-SCs that were stimulated with LTBR-agonist (or isotype control). Right: normalized quantification of CD8^+^ CXCR3^+^ T cells that migrated to the lower transwell chamber. Each dot represents one technical replicate. **c,** Top: experiment to ablate *Ltbr* mosaically and specifically in skin epithelial progenitors by delivering a lentivirus containing *Cre* and *sgLtbr* into the amniotic sacs of E9.5 *R26-LSL-Cas9-P2A-EGFP* pups. Bottom: representative FACS plot and quantification LTBR mean fluorescence intensity (MFI) from EGFP^+^ and EGFP^neg^ uHF-SCs. **d,** qPCR analysis of LTB-LTBR signalling-sensitive transcripts in EGFP-sgLtbr^+^ (red dots) and EGFP^neg^ (white dots) uHF-SCs (top) and IFE-SCs (bottom). **e,** Representative whole mount immunofluorescence and quantification of CD8^+^ T cells in proximity of EGFP^+^ (LTBR-deficient) or EGFP^neg^ (LTBR wild type) HFs. **f,** Schematic shows experiment to ablate all LTB^+^ cells. Following tamoxifen to CreER, *Ltb-CreER; R26-fl-Stop-fl-DTA* targeted (orange) *or R26-fl-Stop-fl-DTA* control (white) mice, animals were colonized with *S. epidermidis* over the course of 6 days. At two weeks, mouse back skin was then swabbed, and bacterial colonies were counted. Right, bar plot of showing number of bacterial colonies obtained from *Ltb* cell-deficient vs littermate control skins. **g,** Experiment to ablate *Ltbr* specifically in skin epithelial SCs and then test the effect on microbial colonization in the skin. Graph shows that bacterial colony numbers correlate with the percentage of *Ltbr-* null mosaicism as quantified by FACS analyses of EGFP^+^ uHF-SCs. Data are representative of two (panel g), three (panel c, d, e) or four (panel b, f) independent experiments and, except in b, each circle or line (in d and e) represents one mouse. Data are analyzed by: Wald test (DESeq2) (panel a); ordinary one-way ANOVA with Tukey’s multiple comparison post-test (b); unpaired two-tailed Student’s *t-*test (c, f); paired two-tailed Student’s *t-*test (d); Wilcoxon matched-pairs signed rank test (e); simple linear regression (g). *p*-values or R squared are indicated in each figure and, when indicated, data are represented as mean with SEM. Further details on statistics and reproducibility in Methods. See Extended Data Fig. 3 and 4 for additional supporting experiments.

To test whether direct activation of LTBR on uHF-SCs caused their upregulation of lymphocyte-chemotactic factors, we performed Boyden Chamber studies *in vitro*, and found that CD4^+^ and CD8^+^ lymphocytes migrated significantly more towards conditioned media collected from uHF-SCs exposed to LTBR-agonist compared to isotype control (**Fig. 3b** and **Extended Data Fig. 4b**). Moreover, anti-CXCL10 blocking antibodies abolished this effect, as seen by reduced migration of CD8^+^ T cells that naturally express the cognate receptor for CXCL10 (CXCR3) (**Fig. 3b**). Notably, the T cell migration response was strongest when conditioned media was from LTBR-agonist-treated uHF-SCs than IFE-SCs or bulge SCs (**Extended Data Fig. 4c**).

These data suggested that epithelial SCs from the skin microenvironment where LTB is enriched are specifically wired to respond to this ligand. The results further indicated that activation of LTBR in uHF-SCs drives a molecular program that orchestrates the creation of an immune-competent niche around HF orifices. In line with this hypothesis, the uHF (LRIG1^+^) showed the closest proximity to CD4^+^ and CD8^+^ T cells and the highest expression of *Cxcl10* and *Cxcl16* compared to the bulge (KRT24^+^) or IFE regions (**Extended Data Fig. 4d-e**).

To test whether LTBR signalling in uHF-SCs is required to recruit and/or maintain lymphocytes around the upper hair follicle *in vivo*, we generated mice where *Ltbr* was mosaically ablated specifically in the skin epithelial SCs. Leveraging the power of *in utero* (E9.5) lentiviral delivery to selectively and stably transduce unspecified skin epithelial progenitors^39^, we mosaically transduced progenitors of *R26-LSL-Cas9-P2A-EGFP* pups by using low titre lentivirus expressing *Ltbr-*targeting small guide RNAs (sgRNAs) and CRE recombinase. This resulted in some skin epithelial SCs expressing EGFP but lacking LTBR (EGFP^+^ LTBR^neg^), while others remained wild type (EGFP^neg^ and LTBR^+^) as validated by flow cytometry (**Fig. 3c**).

To probe the consequences of silencing LTBR in uHF-SCs, we sorted EGFP^+^ and EGFP^neg^ uHF-SCs and observed selective downregulation of *Cxcl10, Cxcl16* and *Nffib2* in EGFP^+^ SCs. Of note, the same analysis carried out on EGFP^+^ and EGFP^neg^ IFE-SCs revealed no significant differences in chemokine expression upon *Ltbr* abrogation (**Fig. 3d**). Thus, the transcriptional effects of LTBR loss on skin epithelial SCs correlated strongly with the status of LTB ligand within the particular SC niche. A notable physiological consequence of these transcriptional changes was the decrease in CD8^+^ lymphocyte recruitment to *Ltbr-*targeted hair follicles (**Fig. 3e**).

When the hair protrudes through the skin surface, the opening offers a hospitable microenvironment for bacterial colonization. As a consequence, the uHF region hosts the highest microbial load in the skin, and several antimicrobial peptides, including defensins, are highly enriched in the uHF-SCs that are tasked with replenishing and rejuvenating the epithelium at these orifices^5–7,40,41^. Recent studies have also unveiled the uHF as the site where adaptive immune responses are mounted upon colonization with *Staphylococcus epidermidis,* a commensal species for human skin^38,42,43^. We therefore posited that LTB-LTBR signalling in the uHF-SCs may be necessary to craft the requisite immune niche that can contain skin bacterial colonization.

To test this hypothesis, we depleted the skin of LTB-expressing immune cells by crossing *Ltb-EGFP-CreER* and *R26-LSL-DTA* mice and then treated topically with tamoxifen during the second resting phase (telogen) of the hair cycle. Two weeks later, skin lacking LTB^+^ cells was colonized with *S.epidermidis,* and bacterial load was tested 1-2 weeks later. As posited, ablation of LTB^+^ cells resulted in a striking failure to contain bacterial colonization (**Fig. 3f** and **Extended Data Fig. 4f**).

To determine whether this effect is rooted in the loss of LTB-LTBR signalling within the uHF-SCs, we again interrogated bacterial load, but this time on mice whose skin epithelial SCs were mosaically ablated of *Ltbr*. Mirroring the effect caused by ablation of LTB-expressing immune cells, a positive correlation was observed between the percentage of skin SCs lacking LTBR and the number of bacterial clones obtained from culturing microbes that had colonized the skin (**Fig. 3g**).

Altogether, our findings unearthed an intimate crosstalk between uHF-SCs and their interacting immune cells, centering on LTB-LTBR signalling to the stem cells. Moreover, this uHF niche-specific crosstalk governed bacterial colonization at the HF orifice.

### Autoregulatory control of DETC abundance in the IFE niche

LAG-3 is known to be elevated in lymphocytes that are stimulated by immunization, infection and/or cancers. There, LAG-3 has been widely studied for its role as an immune checkpoint that restrains immune responses after a challenge, maintains tolerance and/or prevents tumour clearance as an immune-escape mechanism^44–46^. Although constitutive expression of LAG-3 by intraepithelial lymphocytes at barrier sites has been reported^47^, whether and how it might function remained unexplored. We were intrigued to find that in steady state mouse skin, LAG-3 was almost exclusively expressed by IFE-DETCs compared to other lymphocyte populations, including uHF-DETCs (**Fig. 2e** and **Fig. 4a**).

**Figure 4.**
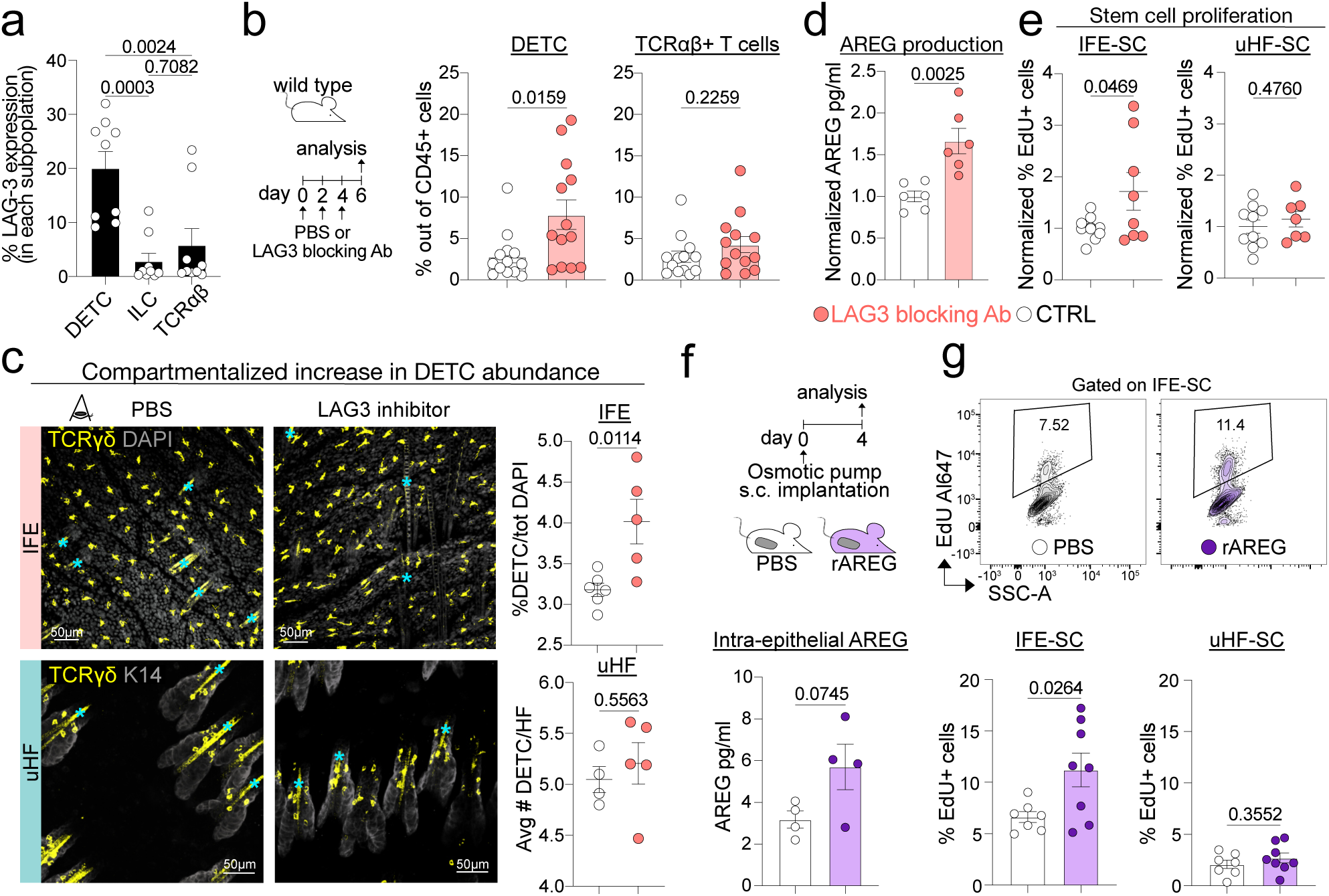
LAG-3 and AREG expression by IFE-DETC converge to govern epidermal homeostasis. **a**, Percentage of LAG-3^+^ cells gated on DETC, ILC or TCRαβ^+^ T cells in wild type back skin as determined by FACS. **b,** Left: strategy to block LAG-3 *in vivo*. Right: quantifications reveal that DETCs but not TCRαβ^+^ T cells increase in numbers in the skin when anti-LAG-3 antibodies are delivered intraperitoneally to mice. **c,** Left: representative max projections of whole mount immunofluorescence of IFE and uHF compartments showing that the increase in DETC (in yellow) resulting from anti-LAG-3 blocking antibodies occurs largely in the IFE-SC and not the uHF-SC niche. Scale bar 50μm. * denote autofluorescence. **d,** Normalized concentration of AREG determined by ELISA shows elevated AREG in the intraepithelial tissue, consistent with the elevation in DETCs caused by LAG-3 inhibition. **e,** Normalized percentage of EdU^+^ SCs determined by FACS after a 3-hour pulse in mice injected with either PBS or anti-LAG-3 blocking antibodies. **f,** Top: strategy to deliver recombinant AREG (rAREG) systemically in vivo by osmotic pump implantation; bottom: concentration of AREG determined by ELISA in the intraepithelial tissue of mice implanted with pumps containing either rAREG or PBS as a control. **g,** Percentage of EdU^+^ SCs determined by FACS after a 4-hour pulse in mice implanted with pumps containing either rAREG or PBS as a control. Data are representative of two (panel f), three (panel c, d, g), five (panel a, e) or seven (panel b) independent experiments and each circle represents one mouse. Data in b, c, d, e, f and g are analyzed by unpaired two-tailed Student’s *t-*test. Data in panel a are analyzed by ordinary one-way ANOVA with Tukey’s multiple comparison post-test. *p-*values are indicated in each figure and data are represented as mean with SEM. Further details on statistics and reproducibility in Methods. See Extended Data Fig. 5 and 6 for additional supporting experiments.

Among reported LAG-3 interacting proteins, Galectin-3 and MHC-II have been the most extensively documented^29,30^. Galectin-3 transcripts (*Lgals3*) were expressed throughout the skin including the epithelium, and while MHC-II (*H2ab1*) was expressed most highly in antigen presenting cells (such as myeloid cells in dermis and Langerhans cells in the epidermis), it was also detected in a subset of epithelial cells (**Extended Data Fig. 5a**). Interestingly, among SCs, expression of LAG-3 interacting proteins was considerably higher in IFE-SCs, mirroring LAG-3 expression by IFE-DETCs (**Extended Data Fig. 5b**).

Seeking a homeostatic role for LAG-3, we used anti-LAG3 blocking antibodies, first substantiating their efficacy by staining activated CD44^+^ CD8^+^ splenocytes with a fluorophore-conjugated antibody targeting the same epitope (**Extended Data Fig. 5c**). We then treated wild type mice with three intra-peritoneal injections of anti-LAG-3 blocking antibodies over the course of one week (**Fig. 4b**). LAG-3 inhibition significantly and selectively increased the proportion of DETCs relative to the total CD45^+^ immune cell population in homeostatic skin (**Fig. 4b**). Moreover, as quantified by immunofluorescence analysis, the expansion of the TCRγδ^+^ T cells in the tissue was restricted to the IFE compared to uHF niches, in line with the enriched expression of LAG-3 by IFE-DETCs (**Fig. 4c**). Together, these results suggested that steady state expression of LAG-3 by epidermal DETCs acts as a short-range brake to control the DETC population size within the IFE-SC niche.

### AREG is an essential modulator of epidermal homeostasis in the IFE-SC niche

To understand how alterations in DETC numbers might affect the skin epidermis, we were intrigued by the expression of amphiregulin by the IFE-DETCs (**Fig. 2**), as it is known to bind to its cognate tyrosine kinase receptor, epidermal growth factor receptor (EGFR), where it can trigger RAS/MAPK signalling. Indeed, the increase in IFE-DETCs upon LAG-3 inhibition was accompanied by an elevation in AREG, as measured by ELISA of the intraepithelial compartment (**Fig. 4d**). Although EGFR is expressed throughout the skin epithelium and skin expresses other EGFR ligands (including EGF, epiregulin and TGFα) (**Extended Data Fig. 6a**), particularly upon injury^27,48,49^, our results hinted that AREG-EGFR mediated immune:stem cell signalling might be critical in governing adult epidermal homeostasis.

In line with a putative regulatory circuit linking DETC numbers to epidermal growth control, LAG-3 inhibition significantly increased *Ki67* expression, a marker of actively proliferating cells, in the intraepithelial compartment (**Extended Data Fig. 5d**). Additionally, a 3-hour pulse with the thymidine analogue 5-ethynyl-2ʹ-deoxyuridine (EdU) following LAG-3 inhibition resulted in enhanced EdU incorporation specifically by the IFE-SCs and not uHF-SCs, mirroring the selective expansion of DETC within the IFE microenvironment upon LAG-3 blockade (**Fig. 4e**).

Linking this proliferative response to the ability of DETCs to produce amphiregulin, we first verified that in agreement with prior *in vitro* studies on human keratinocytes,^50^ mouse IFE-keratinocytes proliferate in response to recombinant AREG (rAREG) (**Extended Data Fig. 6b**). Additionally, when we used osmotic pump delivery to the circulation to elevate intraepithelial AREG levels in the skin to those similar to LAG-3 inhibition, this was sufficient to selectively stimulate proliferation within the IFE (**Fig. 4f-g**).

Finally, to determine whether AREG produced specifically by the IFE DETCs is key in balancing growth and differentiation in the homeostatic epidermis, we generated *Areg fl/fl* x *TCRd-CreER* mice (*Areg* cKO) and compared them to littermate *Areg fl/fl* mice or *Areg* cKO that had not been treated with tamoxifen (CTRL) (**Extended Data Fig. 6c-d**). Even though not complete, the reduction in AREG expression by the DETCs was sufficient to dampen IFE-SC proliferation (**Extended Data Fig. 6e**). By contrast and consistent with the compartmentalization of AREG production by IFE-DETC, proliferation was not affected in the uHF (**Extended Data Fig. 6e**). Together, these results suggested that tonic production of AREG by IFE-DETCs functions in the regulation and maintenance of homeostatic epidermal turnover, while LAG-3 keeps this program in check by regulating the IFE-DETC population size.

### Compensatory maintenance of IFE-SC:immune niche programs

Given the importance of the LAG3-AREG circuit in avoiding epidermal immune cell expansion and over-proliferative stem cell responses, we wondered whether back-up systems may have evolved to preserve this essential regulatory mechanism. As shown in **Fig. 2f-g**, and consistent with a recent multiorgan analysis of TCRγδ T cells^51^, AREG production correlated with DETC colonization of the intra-epidermal niche. To determine whether the epidermal microenvironment is sufficient to drive the acquisition of this immune:stem cell signalling program, we tested whether, in the absence of DETCs, other lymphocytes might be able to take over the vacant IFE niche and acquire *Areg* expression.

In *Areg* cKO mice, DETCs were still present in the IFE niche (**Extended Data Fig. 6d**), and no signs of immune cell compensatory mechanisms were noted. By contrast, when DETCs were absent, as they are in a strain of mice mutant for the *Skint1* gene, the Vγ5^neg^ TCRγδ T cells that are normally found in the dermis, took the place of DETCs within the IFE niche (**Fig. 5a-b**)^26,35^. Intriguingly, while these dermal T cells did not express *Areg* in wild-type skin homeostasis (**Fig. 5c**), in their new residence within the DETC-deficient IFE-SC niche, they not only expressed *Areg* but did so at comparable levels to wild-type DETCs (**Fig. 5c** and **Extended Data Fig. 7a-b**).

**Figure 5.**
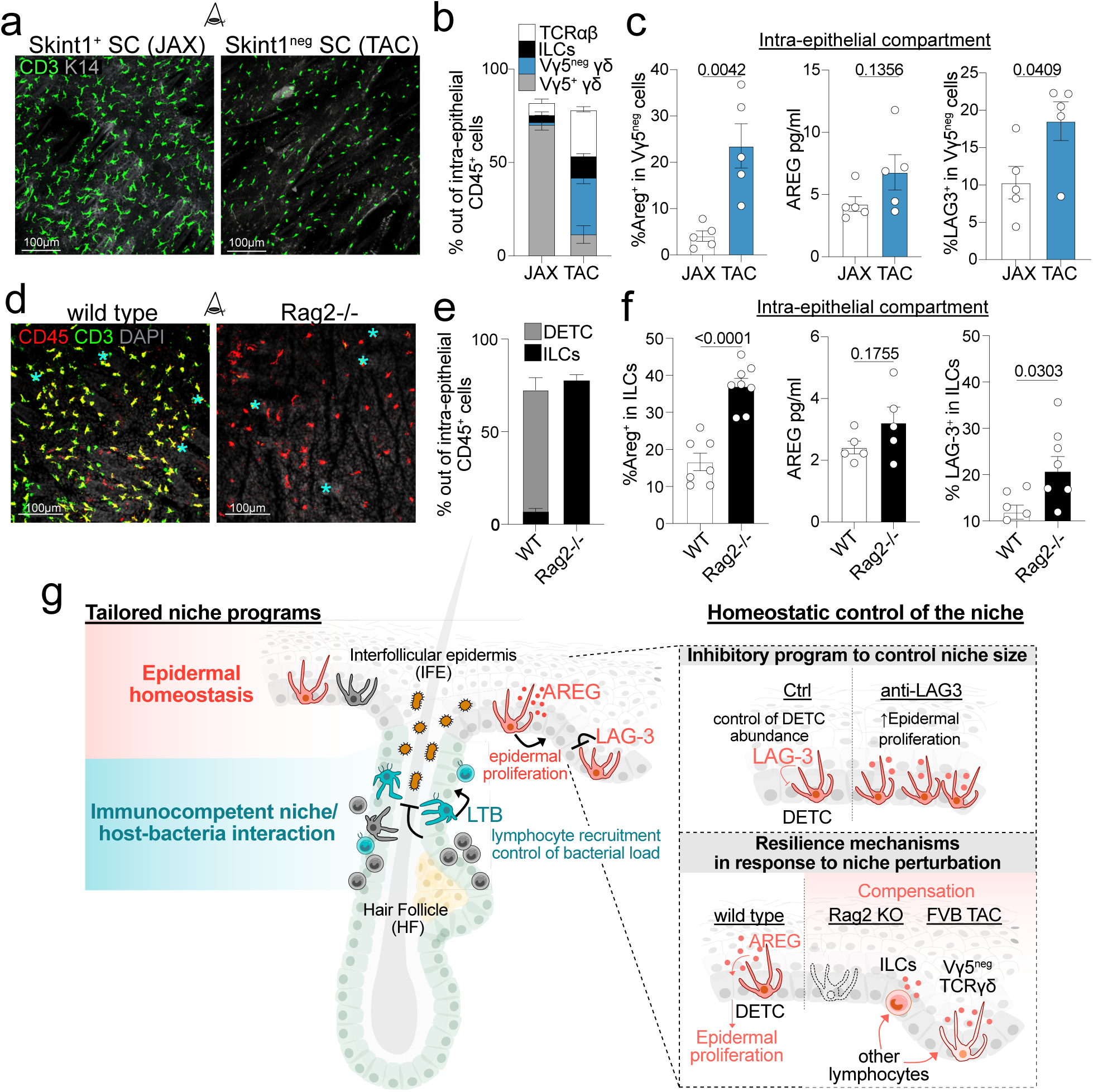
When DETCs are absent, IFE is colonized by and drives new lymphocyte immigrants to express AREG and LAG-3. a, Representative whole mount max projections of the IFE compartment confirms that immigrant T cells (CD3+) occupy the IFE in FVB-TAC (mutant for DETC-cognate ligand, *Skint1*). FVB-JAX mice are shown as WT controls. Scale bar 100μm. **b,** Bar plot showing relative proportions of Vg5+, Vg5neg, TCRab+ T cells and ILCs in the intraepithelial compartment of FVB JAX or TAC mice. **c,** Left: percentage of AREG+ Vg5neg TCRgd T cells detected by FACS in the intraepithelial compartment of FVB JAX or TAC mice. Middle: ELISA showing comparable AREG levels in the intraepithelial fractions from FVB JAX and FVB TAC mice. Right: Representative FACS plots and quantifications of percentage of LAG-3+ Vg5neg TCRgd T cells from the intraepithelial fraction of FVB JAX or TAC mice. **d,** Representative whole mount max projections of the IFE compartment from wild type and Rag2-/-mice showing intraepithelial CD45+ immune cells (red) and CD3+ T cells (green). Scale bar 100μm. * denote autofluorescence. **e,** Quantifications reveal that ILCs (CD45+CD90+ TCRgdneg TCRbneg) compensate for the loss of DETCs (CD45+CD90+ TCRgdhigh) when all T cells are ablated. **f,** Left: FACS analyses of the intraepithelial compartment reveal that ILCs become AREG+ when they occupy the IFE-SC niche vacated by DETC loss. Middle: ELISA reveals that the level of AREG in the intraepithelial fraction is largely independent of the immune resident in the IFE niche. Right: Representative FACS plots and quantifications of percentage of LAG-3+ ILCs from the intraepithelial fraction of wild type (WT) and *Rag2-/-* mice. **g,** Schematic showing immune adaptation within distinct SC niches and homeostatic mechanisms to keep niche size and activity in check. Data are representative of two (panel b, c, e) or three (f) independent experiments, and each circle represents one mouse. Data in c and f are analysed by unpaired two-tailed Student’s *t-*test. *p-*values are indicated in each figure and data are represented as mean with SEM. Further details on statistics and reproducibility in Methods. See Extended Data Fig. 7 for additional supporting experiments.

Probing more deeply into this possible immune niche adaptation within the IFE, we focused on *Rag2*-/-mice, where all T cells including TCRγδ^+^ DETCs are missing. In this case, intraepithelial innate lymphoid cells (ILCs), which are normally found within the uHF and around sebaceous glands^18^, had taken up residency in the IFE, resulting in an expansion in the overall ILC population within the skin (**Fig. 5d-e** and **Extended Data Fig. 7c**)^52,53^. Strikingly, these intraepithelial immigrants upregulated AREG production over that which they made in their WT residences (**Fig. 5f**). Moreover, and intriguingly, the overall amphiregulin transcript expression and protein production in the intraepithelial compartment remained comparable between WT and Rag2-/-skins (**Fig. 5f** and **Extended Data Fig. 7d**). Analogous to AREG, LAG-3 was also markedly upregulated in Vγ5^neg^ TCRγδ T cells and ILCs that colonized the epidermis when DETCs were ablated in FVB TAC and *Rag2-/-* mice, respectively (**Fig. 5c-f**).

Overall, these results provide compelling evidence that whether by DETCs, dermal TCRγδ^+^ or dermal/sebaceous gland ILCs, LAG-3 expression and AREG production is not predetermined, but rather dictated by their colonization within the IFE-SC niche. The ability of the IFE-SC niche microenvironment to fine tune AREG levels further underscores the importance of AREG in controlling homeostatic equilibrium within the epidermis.

## Discussion

Homeostatic stem cell niche programs in tissues rely on the maintenance of a delicate and dynamic equilibrium. Signals from the niche, as well as its composition and critical mass must be tightly coordinated to avoid downstream perturbations that could result in over or underperformance, thereby compromising tissue fitness. For the skin, this is particularly important, as its interfollicular epidermis must continuously replenish the body’s surface barrier while its upper hair follicle must control microbial colonization at the orifice to the outside world.

The stem cell niches dedicated to replenishing and rejuvenating the IFE and uHF must be finely tuned to perform their distinct tasks, but how this happens has long been obscure. Here, using the cutting-edge uLIPSTIC technology to monitor cell-cell contacts, we were able to biotinylate, isolate and characterize unique cell populations that directly interact with the HF stem cell compartments of the skin. By comparing biotinylated versus unlabelled versions of cells that localized to both IFE and uHF SC niches, we discovered that the same cells, in this case Vγ5Vδ1^+^ T lymphocytes (DETCs), undergo regional imprinting within these distinct stem cell niches. In doing so, they tune and tailor their effector functions depending on the interacting stem cell population and its tissue requirements.

We found that in the uHF, DETCs (as well as other niche lymphocytes) express LTB, which triggers immune-SC crosstalk with the LTB receptor-positive uHF-SCs. As we learned, this crosstalk is direct and functions critically in skin homeostasis, by prompting the uHF-SCs to express a cohort of chemokines that in turn attract and recruit other immune cells that participate in protecting the HF orifice against excessive microbial load^18^. Here, our genetic studies further revealed that if the stem cells cannot express the LTB receptor, they fail to generate the requisite chemokines that bring additional protective immune cells to the niche. The consequences are substantial, as illustrated by the marked increase in microbial burden upon commensal colonization.

We unearthed strikingly different, but similarly surprising, roles for the DETCs in the homeostatic epidermis. While EGFR signalling has long been known to function critically in controlling epidermal growth and differentiation, a role for DETCs in this process has been largely overlooked. As we showed, not only is the immune niche within the epidermis responsible for producing AREG and regulating stem cell proliferation, but in addition, the magnitude of this proliferative effect is directly controlled by DETC density within the IFE niche. This turned out to be conferred by DETC expression of LAG-3, acting as a brake to keep DETC niche levels in check (**Fig. 5g**).

Finally, and as importantly, our study exposed resilience mechanisms to ensure that the critical immune:epithelial crosstalk between skin stem cells and their niches is maintained for the sake of tissue fitness. Thus, when DETC vacancies were generated within the IFE, at least two different dermal immune cell types—ILCs and γ5^neg^ γδT cells—moved into the IFE. There, they co-adapted within their new tissue microenvironment to acquire the key factors of the IFE-DETC transcriptome that are needed to preserve the balance of AREG/LAG3-producing immune cells and IFE-SCs.

Overall, our studies on the SC: immune cell interactions within two distinct stem cell niches place the stem cells at the helm of weaving their niches and the immune cells at the helm of maintaining them. In this regard, it is interesting to consider a similar phenomenon may occur in the bone marrow where distinct subpopulations of leptin receptor^+^ (LEPR) mesenchymal niche cells populate the hematopoietic SC and lymphoid progenitor niches. Analogous to the DETCs in the IFE and uHF skin stem cells niches, these two LEPR^+^ populations are ontogenically related^8^. In the skin, SC niche adaptation also occurs, this time as a fine tuning of effector functions within the DETCs. Remarkably, however, as we discovered, SC-specific adaptation was not restricted to ontogenically related immune cells, but rather controlled by the niche microenvironment and its vacancies.

In closing, we have discovered that immune cell occupants of SC niches are dynamic, pliable and can adapt their program of gene expression to preserve the necessary immune-SC crosstalk to ensure epithelial fitness. The existence of compensatory mechanisms to preserve this crosstalk are particularly relevant in reconciling the fact that human skin does not harbour DETCs but rather is populated by other tissue resident T lymphocytes. Our results portray a model where co-adaptation between SCs and their immune niches is at play in different tissue microenvironments and that quorum sensing-like mechanisms, the details still unfolding, have evolved to maintain critical niche programs at homeostatic levels in both steady state and in response to niche perturbation (**Fig. 5g**).

## Data Availability

All data supporting the findings of this study are available within the Article. Single-cell and bulk sequencing data generated within this study have been deposited and will be made available upon publication at the Gene Expression Omnibus (GEO) under the accession code GSE314678 (bulk RNA-seq) and GSE314679 (single-cell RNAseq).

## Code Availability

Custom code for scRNA-seq for this study will be deposited upon publication in GitHub.

## Acknowledgements

The authors thank E. Wong, M. Nikolova, J. Racelis, L. Hidalgo, I. Crawley for technical assistance; J. Almagro, M.D. Abdusselamoglu, C. Cowley, Y. Yang, N. Guzzi, M. Kudelka, I. Izgi, C. Xu, M. Schernthanner, A. Gola, N. Alexander, J. Castellanos, W. Song, R. Niec, G. Monasterio and E.J. Villablanca for discussions and protocols; The Rockefeller University FACS facility (S. Mazel, director) and the Weill Cornell Medicine and Rockefeller University Genomics Resources Core Facilities. E.F., D.M. and G.D.V. are Howard Hughes Medical Investigators. S.M.P. was the recipient of the Nicholson Postdoctoral Fellowship, the CRI/Carson postdoctoral fellowship (CRI4498) and the NIH NIAMS K99/R00 Pathway to Independence Award (5K99AR084573-01). S.M.S is supported by a Ruth L. Kirschstein Predoctoral Individual National Research Service Award (F31AR083275) and a recipient of the National Institutes of Health (NIH) Clinical and Translational Science Award (CTSA) through Rockefeller University. A.R.B. was the recipient of a Damon Runyon Postdoctoral Fellowship and is now a Hanna H. Gray Fellow of the Howard Hughes Medical Institute. S.Y. was the recipient of an F31 Ruth L. Kirschstein Predoctoral Individual National Research Service fellowship from the National Cancer Institute (NCI) and a Pilot Award from the Shapiro-Silverberg Fund at The Rockefeller University. S.A.L. was supported by NIH R01DK110352. G.D.V. was supported by NIH R01AI173086. E.F. is the recipient of a research award from the Stavros Niarchos Foundation Institute for Global Infectious Disease Research and a Discovery Award from the Glenn Foundation. This study was supported by grants to E.F. from the National Institutes of Health (R01-AR050452, R37-AR27883 and R01-AR31737).

## Contributions

S.M.P. and E.F. conceptualized the study, designed the experiments, interpreted the data and wrote the paper. S.M.S. analysed the scRNAseq and bulk RNAseq data. C.B. assisted with the pre/postnatal kinetic analyses and FVB analysis. V.M.O. assisted with the initial anti-LAG3 experiments. S.Y. helped performing the uLIPSTIC scRNAseq experiment. A.R.B. and S. N-H. helped with the initial uLIPSTIC experiments. S.A.L. generated the Areg fl/fl mouse strain. D.M. and G.D.V. contributed germ-free animals, uLIPSTIC mice and expertise and scientific input on the study. All authors provided input on the final manuscript.

## Extended Data Figures

**Extended Data Figure 1.**
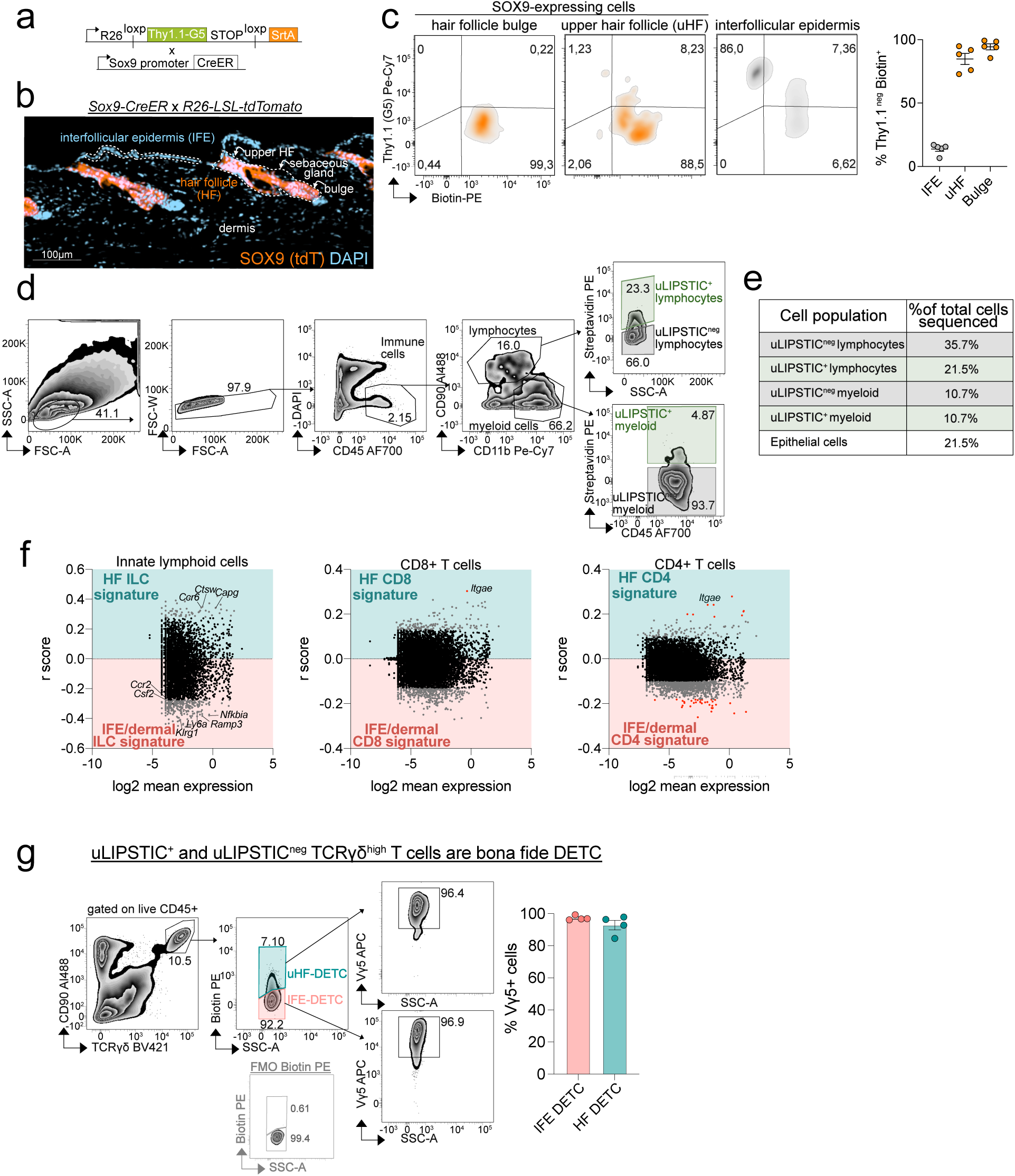
Validation of uLIPSTIC x Sox9-CreER mice as a tool to label the HF-SC immune niche. **a,** Schematic showing the genotype and crossing strategies to generate the hair follicle interactome reporter mice (*uLIPSTIC x Sox9-CreER* mice). **b,** Representative immunofluorescence from a sagittal skin section showing the distribution of SOX9 expression (i.e. restricted to the HF-SC compartment) from *Sox9-CreER x R26-tdTomato* second telogen mice. **c**, Representative FACS plots (left) and quantification (right) of uLIPSTIC donor cells defined as Thy1.1^neg^ Biotin^+^ among bulge-SC, uHF-SC or IFE-SC. Each dot represents one mouse from four independent experiments. **d**, Representative FACS plots showing the gating strategy to isolate uLIPSTIC^+ and neg^ lymphocytes and myeloid cells from *uLIPSTIC-Sox9-CreER* mice. **e**, Comparable number of uLIPSTIC^+ and neg^ cells were sequenced (not representative of the *in vivo* relative proportions). The table shows the relative proportion of each sequenced cell type out of all the cells that were sequenced. Epithelial SC from wild type mice were included to analyse expression of receptors for immune-derived ligands. **f**, volcano plot showing correlation of uLIPSTIC barcode signal and log2 mean gene expression in innate lymphoid cells, CD8^+^ and CD4^+^ T cells from *uLIPSTIC-Sox9CreER* mice sequenced by 10x scRNAseq. **g**, Representative FACS plots (left) and quantification of Vγ5^+^ cells out of Biotin^+^ (uLIPSTIC^+^) and Biotin^neg^ (uLIPSTIC^neg^) TCRγδ^high^ T cells. Each dot represents one mouse.

**Extended Data Figure 2.**
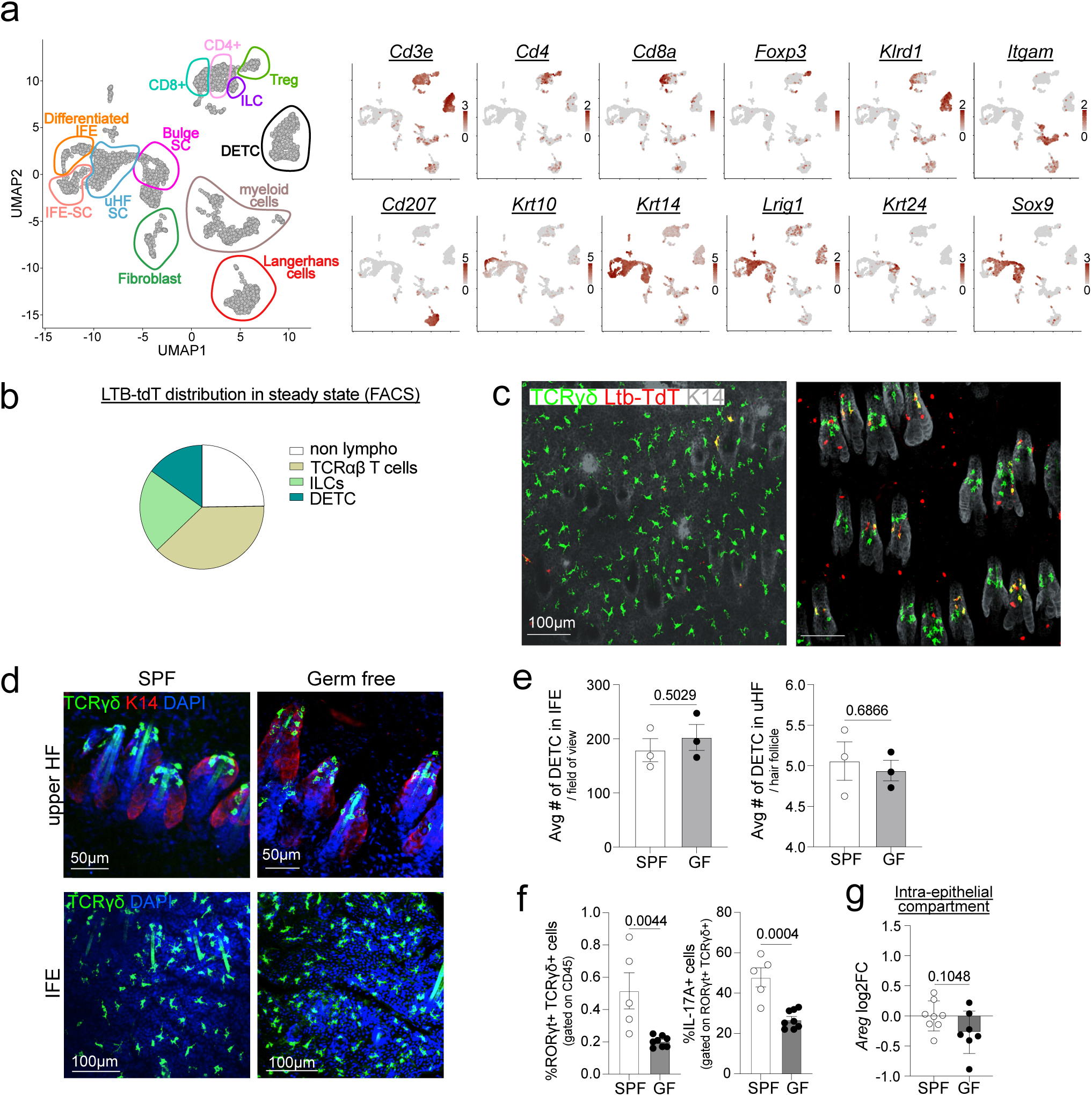
Distribution of LTB^+^ cells in LTB-reporter mice and characterization of the immune niche in germ free mice. **a**, UMAP plots showing expression intensity of genes used to annotate cell types. **b**, Pie chart showing relative proportions of TCRαβ^+^ T cells, ILCs, DETC and other non-lymphocytic cells (i.e. CD90^n^^eg^) out of total LTB-tdT^+^ cells from LTB-reporter mice. **c**, Representative max projections from whole mount immunofluorescence of the IFE and uHF region in *Ltb-EGFP-CreER x R26-LSL-tdTomato*. **d**, Representative max projections of whole mount immunofluorescence of the IFE and uHF compartment of SPF and GF back skin. DETC are identified by anti-TCRγδ antibody staining and shown in green. **e**, quantification of the average number of DETC in the IFE and uHF of SPF and GF back skin. For IFE-DETC, the average number of DETC per field of view is plotted. For the uHF DETC, the average number of DETC per hair follicle is plotted. Each dot represents one mouse. **f**, Bar plots showing the percentage of RORγt^+^ TCRγδ^+^ T cells out of total CD45^+^ cells and IL17A^+^ cells out of RORγt^+^ TCRγδ^+^ T cells in the back skin of specific pathogen free (SPF) and germ-free (GF) mice. Each dot represents one mouse from two independent experiments. **g**, qPCR analysis of *Areg* transcripts in the intraepithelial fraction of SPF and GF mice. Each dot represents one mouse from three independent experiments. Data in e, f and g are analysed by unpaired two-tailed Student’s *t-*test. *p*-values are indicated in each figure and data are represented as mean with SEM.

**Extended Data Figure 3.**
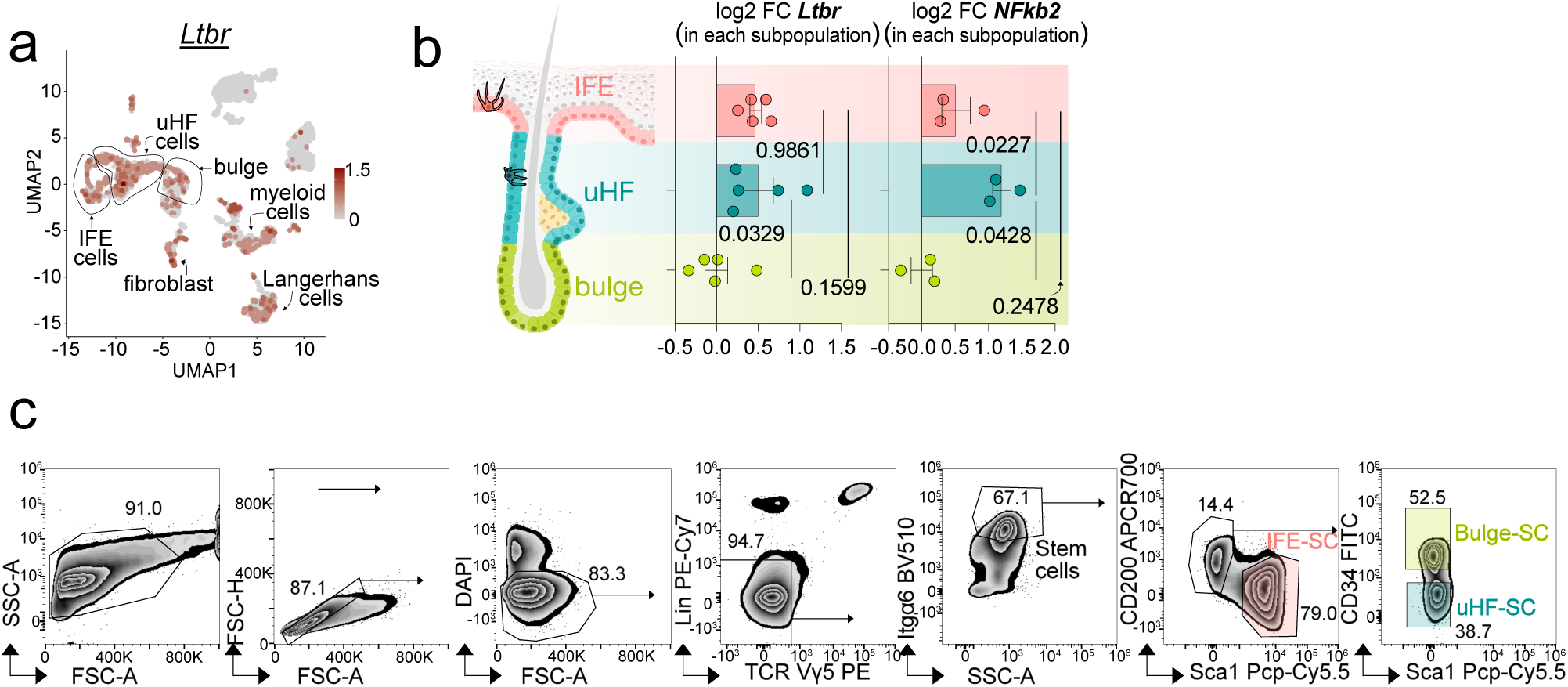
Expression pattern of LTBR and its downstream signalling. **a**, UMAP plots showing expression pattern of *Ltbr* across different cell type from mouse back skin. **b**, qPCR analysis of *Ltbr* and *Nffib2* expression Bulge-SC, uHF-SC and IFE-SC FACS-sorted from wild type mice. Each dot represents one mouse from one-two independent experiments. **c**, Gating strategy used to isolate Bulge-SC, uHF-SC and IFE-SC. Data in b are analysed by ordinary one-way ANOVA with Tukey’s multiple comparison post-test. *p*-values are indicated in each figure and data are represented as mean with SEM.

**Extended Data Figure 4.**
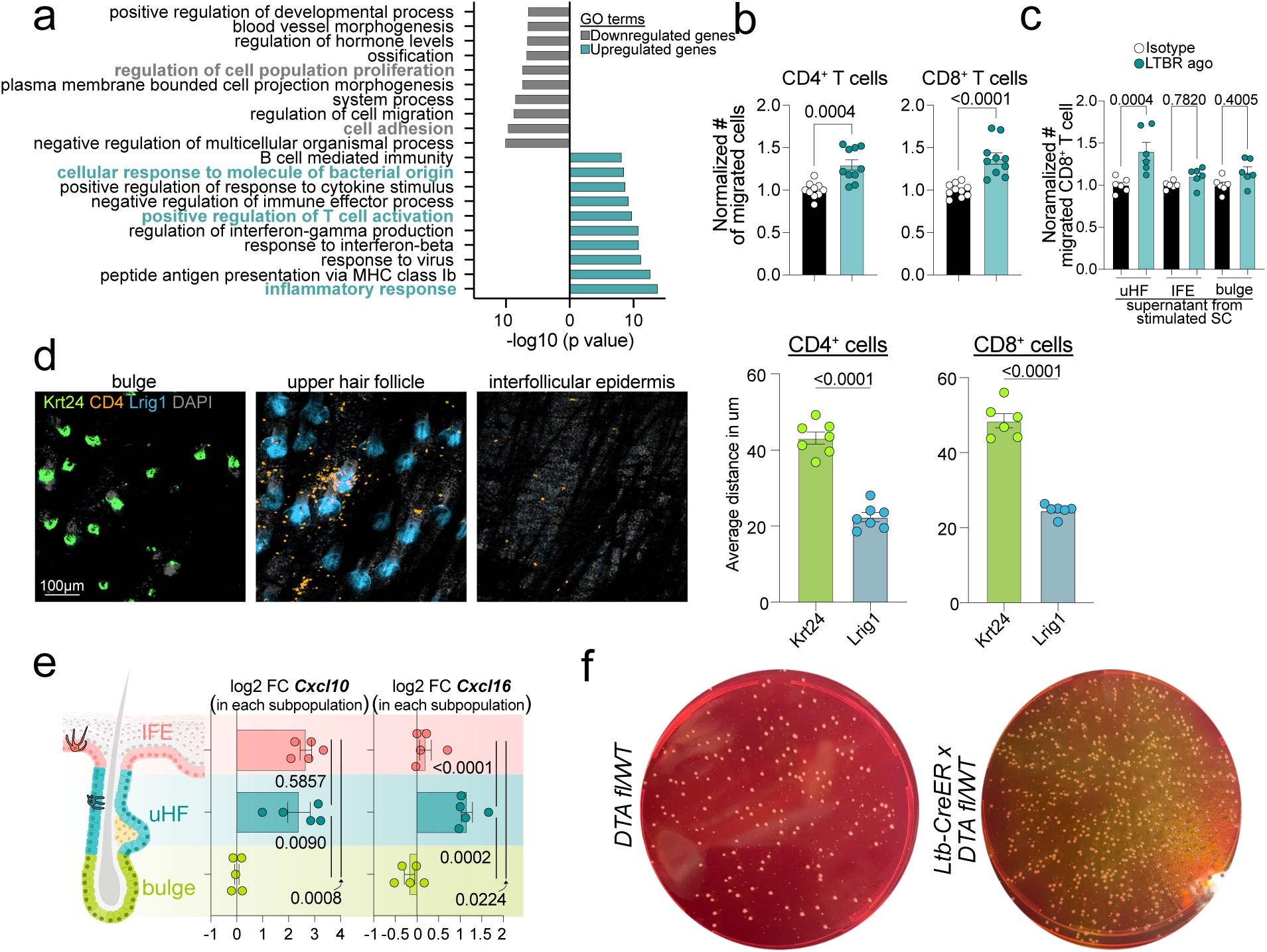
Characterization of LTBR signalling in WT mice treated or not with LTBR agonist. **a**, GeneOntlogy (GO) terms up- and down-regulated in uHF-SC upon in vivo intradermal administration of LTBR-agonist compared to isotype control injected mice. **b**, Normalized quantification of number of CD4^+^ and CD8^+^ T cells migrated through a transwell membrane towards the supernatant collected from uHF-SC treated with LTBR agonist or isotype control. Each dot represents one technical replicate from four independent experiments. **c**, Normalized quantification of number of CD8^+^ T cells migrated through a transwell membrane towards the supernatant collected from Bulge-SC, IFE-SC or uHF-SC treated with LTBR agonist or isotype control. Each dot represents one technical replicate from three independent experiments. **d**, Left: Representative max projections of whole mount immunofluorescence of the bulge, uHF and IFE compartment of wild type mice showing distribution of CD4^+^ cells (in white). Bulge cells are marked by Krt24 in green and upper hair follicle cells are marked by LRIG1 in light blue. Right: Quantification of the average distance of CD4^+^ and CD8^+^ T cells from Krt24 and LRIG1 quantified upon surface rendering, *n*=2 mice. **e**, qPCR analysis of *Cxcl10* and *Cxcl16* expression Bulge-SC, uHF-SC and IFE-SC FACS-sorted from wild type mice. Each dot represents one mouse from two independent experiments. **f**, Representative images of MSA plates showing bacterial colonies grown after swabbing the back skin of *Ltb-CreER x R26-DTA* and *R26-DTA* (ctrl) mice associated with *S. epidermidis*. Data in b and d are analysed by unpaired two-tailed Student’s *t-*test. Data in c and e are analysed by ordinary one-way ANOVA with Tukey’s multiple comparison post-test. *p*-values are indicated in each figure and data are represented as mean with SEM.

**Extended Data Figure 5.**
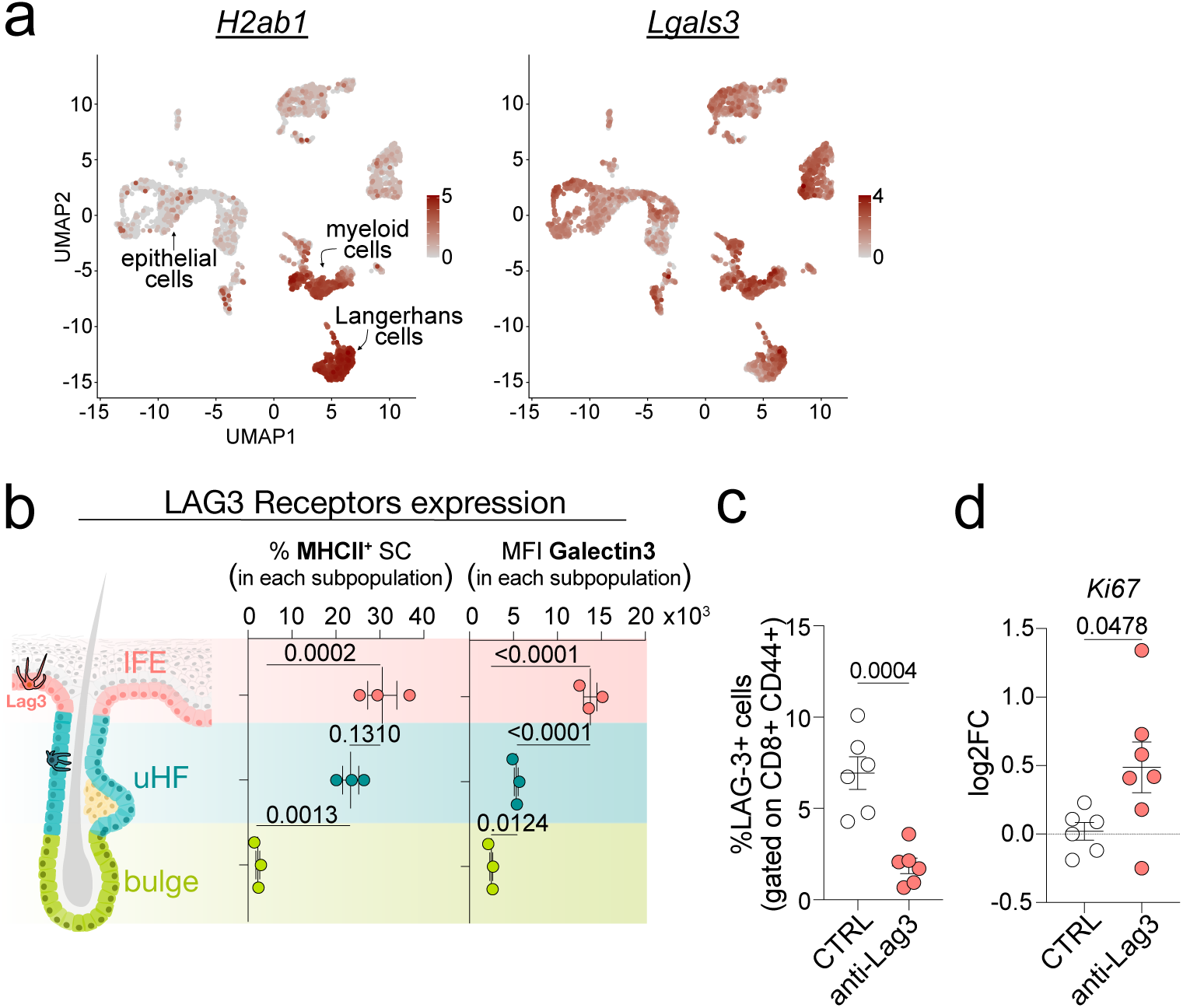
Expression pattern of LAG-3 receptors and characterization of anti-LAG-3 *in vivo* treatment in skin. **a**, UMAP plots showing expression pattern of *H2-ab1* (MHC-II) and *Lgals3* (Galectin-3) across different cell type from mouse back skin. **b**, Quantification of the percentage of MHC-II^+^ and mean fluorescence intensity (MFI) of Galectin-3 in Bulge-SC, uHF-SC and IFE-SC from wild type mice determined by FACS. Each dot represents one mouse. **c**, Percentage of LAG-3^+^ cells gated on CD8^+^ CD44^+^ splenocytes from wild type mice injected with PBS or with anti-LAG3 blocking antibodies. Each dot represents one mouse from two independent experiments. **d**, qPCR analysis of *mKi67* expression in the intraepithelial compartment of wild type mice injected with PBS or with anti-LAG3 blocking antibodies. Each dot represents one mouse from three independent experiments. Data in c and d are analysed by unpaired two-tailed Student’s *t-*test. Data in b are analysed by ordinary one-way ANOVA with Tukey’s multiple comparison post-test. *p*-values are indicated in each figure and data are represented as mean with SEM.

**Extended Data Figure 6.**
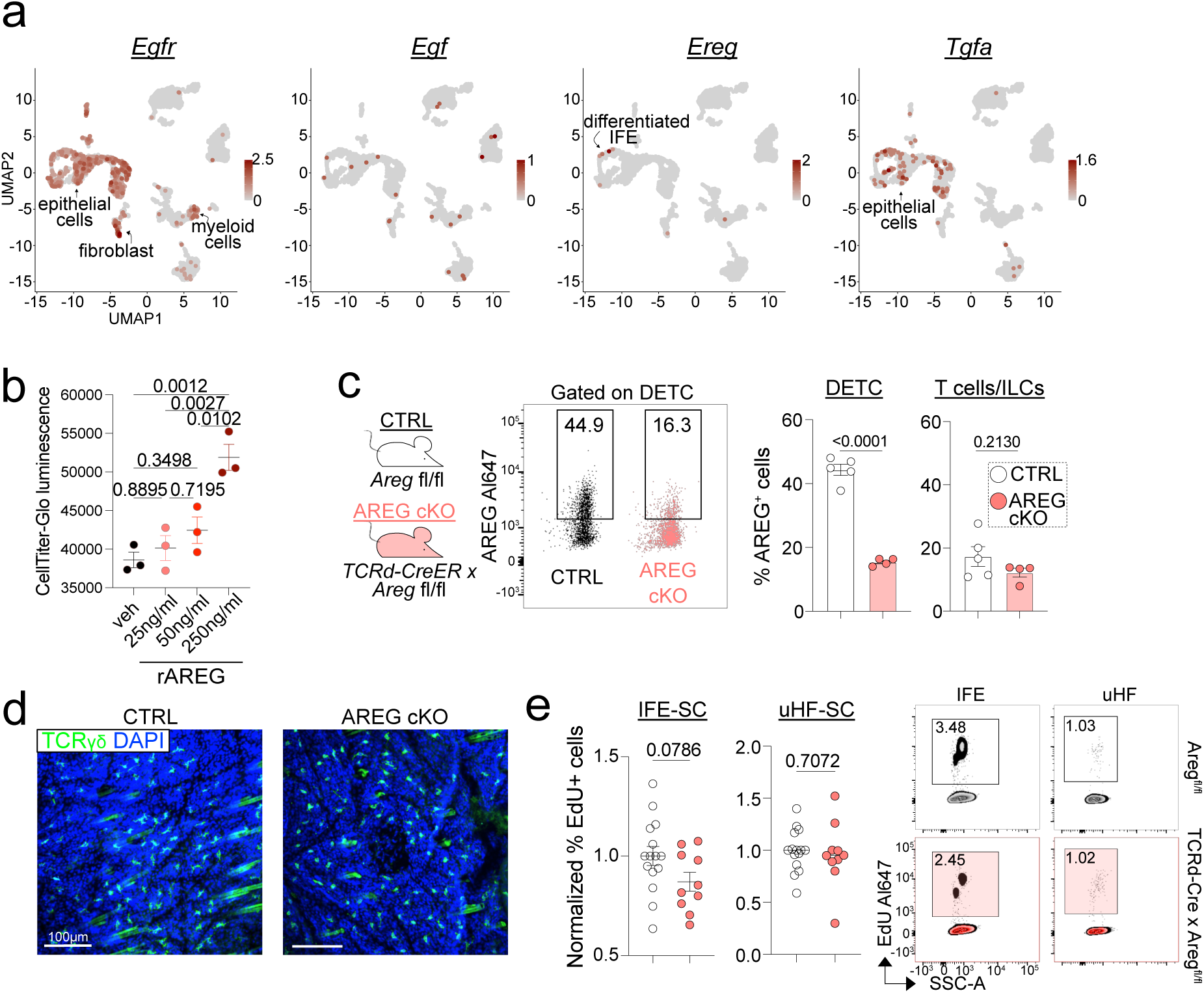
Characterization of the IFE niche in mice with AREG conditional knockout in TCRγδ T cells. **a**, UMAP plots showing *Egfr, Egf, Tgfa* and *Ereg* expression pattern across different cell types from mouse back skin. **b**, Dose-dependent stimulation of IFE-SC proliferation *in vitro* by recombinant AREG. Analysis was by CellTiter-Glo luminescence assay. Each dot represents a technical replicate, and data are representative of two independent experiments. **c**, Conditional ablation of *Areg* in DETCs was achieved by mating *TCRδ-CreER* to Aregfl/fl mice. Analyses of skin immune cells revealed that DETCs, the major source of AREG, show efficient targeting, relative to *TCRδ^neg^* T cells and ILCs, which express much lower AREG than DETCs and are unaffected. Each dot represents one mouse from three independent experiments. Note that AREG loss did not affect DETC numbers. **d**, Representative max projection of whole mount immunofluorescence of the IFE compartment from CTRL and AREG cKO mice showing distribution of DETC (stained with TCRγδ antibodies) in the IFE. **e**, Representative FACS plots (right) and normalized percentage (left) of EdU^+^ cells in IFE-or uHF-SC detected by FACS in CTRL or AREG cKO mice after 4-hour pulse. Each dot represents one mouse from 7 independent experiments. Data in b are analysed by ordinary one-way ANOVA with Tukey’s multiple comparison post-test. Data in c, and e are analysed by unpaired two-tailed Student’s *t-*test. *p*-values are indicated in each figure and data are represented as mean with SEM.

**Extended Data Figure 7.**
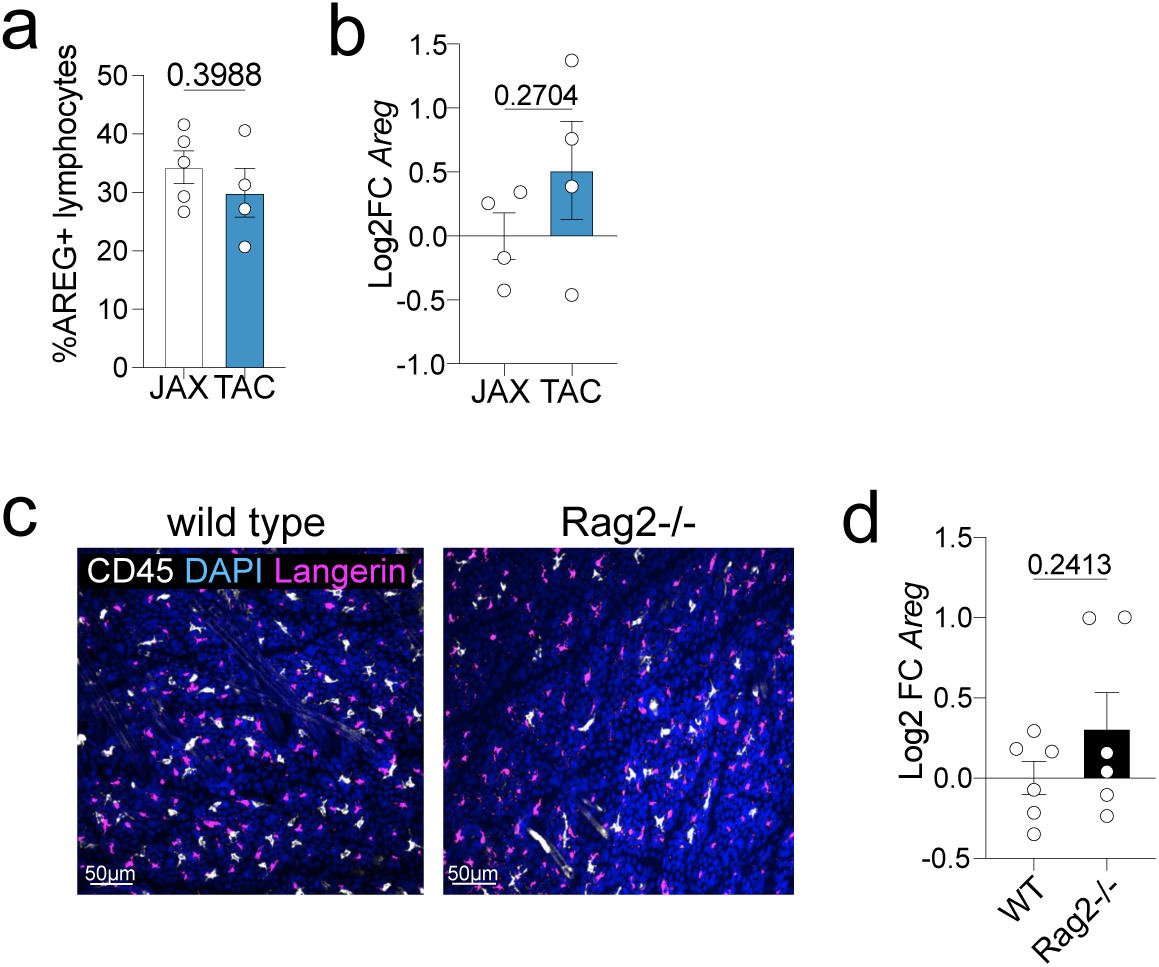
Characterization of the IFE niche in mice whose epidermal stem cell niche is colonized by different lymphocytes. **a**, Bar plots showing the percentage of AREG^+^ intraepithelial lymphocytes (defined as CD90^+^) from FVB TAC and JAX mice. Each dot represents one mouse from two independent experiments. **b**, qPCR analysis of *Areg* transcripts in the intraepithelial fraction of FVB JAX or TAC mice. Each dot represents one mouse from two independent experiments. **c**, Representative max projection of whole mount immunofluorescence of the IFE compartment of wild type and Rag2-/-mice showing distribution of Langerhans cells (defined as CD45^low^ and Langerin^+^) and other immune cells (CD45^+^) populating the IFE. **d**, qPCR analysis of *Areg* transcripts in the intraepithelial fraction of WT or Rag2-/-mice. Each dot represents one mouse from two independent experiments. Data in a, b and d are analysed by unpaired two-tailed Student’s *t-*test. *p*-values are indicated in each figure and data are represented as mean with SEM.

## Methods

### Animals

C57BL/6J wild type mice were purchased from The Jackson Laboratories (strain#000664) and acclimatized in house for at least one week before being used for experiments. uLIPSTIC mice (C57BL/6-Gt(ROSA)26Sortm1(CAG-Thy1,-srta)Vic/J, JAX strain#038221) were obtained from the Victora lab at the Rockefeller University and generated as reported in^1^. Sox9-CreER mice (C57BL/6J background) were provided by Haruhiko Akiyama/Benoit Lab (The University of Texas MD Anderson Cancer Center). Ltb-EGFP-CreER mice (C57BL/6N-Atm1Brd Ltbtm2(EGFP/cre/ERT2)Wtsi/WtsiIeg) were obtained from EMMA, rederived in house and crossed to Rosa26lsl-tdTomato (B6.Cg-Gt(ROSA)26Sortm14(CAG-tdTomato) Hze, JAX strain #007914) or to Rosa26lsl-DTA (B6.129P2-Gt(ROSA)26Sortm1(DTA)Lky/J JAX strain#009669). Tcrd-CreER (B6.129S-Tcrdtm1.1(cre/ERT2)Zhu/J JAX strain#031679) were obtained from the Mucida Lab at the Rockefeller University. Areg fl/fl mice were obtained by Sergio Lira (Icahn School of Medicine, New York) and generated as described in^2^. Rag2 KO animals were originally obtained from The Jackson Laboratories (JAX strain# 008449) and they were bred and maintained in house. Germ-free animals were obtained from the Mucida lab at the Rockefeller University. C57BL/6J wild type mice housed in the same animal facility were used as a control for Rag2-/- and germ-free experiments. FVB/NJ were purchased from the Jackson Laboratories (FVB/NJ, strain# 001800), while FVBn/TAC were purchased from Taconic Biosciences (FVB/NTac). Rosa26-LSL-Cas9-EGFP (B6J.129(B6N)-Gt(ROSA)26Sortm1(CAG-cas9*,-EGFP)Fezh/J, JAX strain#026175) were purchased from JAX and bred in house.

Mice were assigned randomly to experimental groups and age-, sex-matched and littermates were used for each experiment (except for studies with Rag2-/-, germ-free and FVB from TAC and JAX). Both male and female mice were used in each experiment (except for the LIPSTC scRNAseq experiment where only female mice were used). Unless indicated, mice were used during the resting phase of the hair cycle (second or third telogen). Mice were maintained in the Association for Assessment and Accreditation of Laboratory Animal Care. AAALAC-accredited Comparative Bioscience Center of The Rockefeller University (RU), and procedures were performed with Institutional Animal Care and Use Committee (IACUC)-approved protocols. Mice were housed in an environment with controlled temperature and humidity, on 12 h light:dark cycles, and fed with regular rodent chow (5053).

For labelling of CreER reporters, 100-150μl of 20mg/ml of tamoxifen (Sigma Aldrich) resuspended in corn oil (Sigma Aldrich) were administered intraperitoneally to the mice. Depending on the strain 5-7 injections over two weeks were administered and mice were analysed 1-2 weeks after the last tamoxifen injection. For labelling of proliferating cells, 50μg/g of EdU were intraperitoneally injected 4hr before lethal administration of CO2.

### uLIPSTIC substrate administration

Biotin-LPTEG substrate was purchased from LifeTein (biotin–aminohexanoic acid–LPETGS, carboxy-terminal amide, at 95% purity) and resuspended in PBS at 20 mM (stock solutions). Sox9CreER x uIPSTIC mice were injected intraperitoneally with 100μl of stock solution for six times every 20 minute. Mice were sacrificed and the skin was harvested 20-40 minutes after the last injection.

### Whole-mount immunofluoresence

Hairs on mouse back skin were first shaven and then fully removed with hair removal cream (Veet). Skin tissue (approx. 1 cm^2^) was excised with forceps and scissors, and big areas of fat tissue were removed. Tissue was placed on Whatman paper and fixed in 4% paraformaldehyde (Electron Microscopy Science) in PBS for 1 hour at room temperature (RT), followed by extensive washing in PBS. The tissue was further trimmed (approx. 3-4 mm^2^) and placed in 0.3% Triton X100 in PBS (PBST) overnight at RT with horizontal shaking. For immunolabeling, primary antibodies were diluted in blocking buffer (PBS, 5% Normal Donkey Serum, 0.3% Triton X100, 1% bovine serum albumin, 1% fish gelatin) and skin tissues were incubated for 24-48 hours at RT with horizontal agitation. Samples were then extensively washed in PBST and incubated at RT for 24h with secondary antibodies conjugated with Alexa 488, Alexa 647 or RRX (1:400 Life Technologies) and DAPI (0.2 μg/ml) in blocking buffer. Samples were then washed extensively in PBST, quickly dried on a tissue and immersed in clearing solution (RapiClear 1.52, SUNJin Lab). Cleared skin was mounted between a glass slide and a cover slip and placed in the microscope holder for image acquisition. The following primary antibodies were used in the study: chicken anti-keratin 14 (Elaine Fuchs’lab, 1:300); chicken anti-GFP (AVES Lab, 1:500); goat anti-LRIG1 (RnD System, 1:50); rabbit anti-keratin 24 (Elaine Fuchs’lab, 1:500); hamster anti-Vγ5 FITC (536, BD Biosciences, 1:100), hamster anti-TCRγδ Alexa488 (GL3, Biolegend, 1:100); rat anti-CD45 (S18009D, Biolegend, 1:100); rat anti-mouse CD3e (17A2, Biolegend, 1:100); rat anti-CD8 (53-6.7 Biolegend, 1:150); rat anti-CD4 (GK1.5, Biolegend, 1:150), rat anti-Langerin (929F3.01, Novus Biological, 1:300).

### Tissue digestion and single cell suspension isolation

#### Stem cells and intraepithelial fraction isolation

mouse back skin was shaven and excised and big pieces of fat removed with forceps. Dermal tissue was removed by scraping with a dull scalpel and the tissue was placed (epidermis up) onto Trypsin EDTA (Gibco, 0.25% in PBS) for 35-45 minutes at 37°C with horizontal agitation. The tissue was then scraped with a dull scalpel (on the epidermal side) to release the cells in the media. The resulting cell suspension was quenched using FACS buffer (PSB + 5% FBS), transferred to a 50ml conical tube (Falcon), shaken vigorously and sequentially filtered through a 70μm and 40μm filters. Cells were then spun down at 1700 rpm for 5 minutes and the pellet was either directly stained with primary antibodies for flow cytometry, resuspended in RLT buffer (Qiagen) for RNA extraction or processed with Percoll gradient (see below, as in the case of cytokine stimulation for DETC).

#### Immune cell isolation from the whole skin

mouse back skin (approx. 2-3 cm^2^) was excised, and big pieces of fat were removed with forceps. The tissue was placed (dermis down) onto HBSS (Gibco) + collagenase type I (Invitrogen 4mg/ml) at 37°C for 1 hour and 15 minutes with horizontal agitation. Cells released in suspension were collected into a 50ml conical tube and placed on ice (Fraction 1), the remaining tissue was incubated (still dermis down) with Trypsin EDTA for 35 minutes at 37°C. Cells were released in the media by vigorous scraping (on the epidermal side) with a dull scalpel and the resulting cell suspension was quenched with FACS buffer and pooled with the respective Fraction 1. Cells were then sequentially filtered through 70μm and 40μm filters and spun down at 1700 rpm for 5 minutes. The resulting pellet was processed for Percoll gradient enrichment of lymphocytes.

#### Percoll gradient enrichment of immune cells

a 100% solution of Percoll was prepared by combining Percoll (Sigma Aldrich) with 10% of 10x PBS. Two solutions of 40% and 80% Percoll were then prepared by diluting the 100% Percoll in HBSS. Cell pellets were resuspended in 6ml of 40% Percoll, transferred to a 15ml conical tube (Falcon) and 1.8ml of 80% Percoll solution was gently underlaid, creating a two-phase solution. Tubes were centrifuged at 800g, RT for 20 minutes with slow acceleration and no break. The visible ring of cells enriched at the two solutions’ interphase was transferred to another 15ml conical tube, extensively washed with FACS buffer and spun down at 1700rpm for 5 minutes.

#### Splenocytes isolation

spleen was harvested and cleaned form fat with forceps and scissors. The organ was placed on a 70μm cell strainer and smashed using the back side of a 1ml syringe plunger. The filter was extensively washed with FACS buffer and cells were spun down at 1500rpm for 5 minutes before proceeding with primary antibodies incubations.

### Embryonic thymocytes isolation

pregnancy was timed in order to harvest embryos at the desired time point. Pregnant moms were sacrificed and the embryos harvested and placed on cold PBS. Under a dissection scope, embryos were immobilized and the chest area opened with a longitudinal incision. Once localized (above the embryonic heart), the two thymic lobes were collected using curved forceps and placed in cold FACS buffer. Single cells suspension was obtained by smashing the organs on 70μm cell strainers with the back side of a plunger and extensively washing with FACS buffer.

### 10x Single cell RNA sequencing analysis

#### Library preparation

cell suspension from uLIPSTIC mice back skin was stained with fluorescent antibodies and with HTO-streptavidin for detection of the uLIPSTIC signal (streptavidin-PE TotalSeq-C0961 barcoded antibody, Biolegend, 1:1000). Cells were sorted in eppendorf tubes and mixed together according to the ratios shown in Extended Data Fig. 1e. Cells were centrifuged and resuspended in approx. 50 μl of PBS. Cells were counted for viability before being submitted for library preparation with 10X Single cell Chromium system (5’ 10x scRNAseq + TotalSeq C feature barcode + VDJ library prep) according to manufacturer’s instructions (Genomics Core of Rockefeller University). The lybrary was then sequenced on an Illumina Novaseq SP flowcell.

#### Analysis

Raw fastq files were aligned to the mouse genome (mm10, v1.20 from 10X Genomics) and quantified using the ‘count’ function in CellRanger (v6.0.0) with default parameters. The resulting raw counts matrices were then loaded into Seurat (v5.0.1). Raw hashtag oligomer (HTO) counts corresponding to uLIPSTIC signal were normalized per cell as [count_HTO_ / (count_Total RNA_ + count_HTO_)]*10^3^, and RNA expression values normalized using the sctransform function with parameter “flavor=vst1”. The resulting Seurat object was annotated for epithelial, myeloid, and lymphoid compartments as in Extended Data Fig. 2A. For analysis of all immune cells (see **Fig. 1C**), this Seurat object was then subset to exclude epithelial cell spike-in (expressing Krt1, Krt5, Krt10, or Krt14) and Langerhans cells (expressing Cd207), which have been disproportionately enriched for the uLIPSTIC^+^ fraction (over the uLIPSTIC^neg^ fraction) and hence have been removed from the analysis to avoid misleading results. For analyses of individual cell types, each cell type was subset from this Seurat object into a separate object and re-normalized as above.

For correlation analyses between uLIPSTIC signal and gene expression, normalized HTO and normalized RNA expression values from Seurat objects (individually subset for each cell type, as above) were assessed for Spearman correlation, and associated false-discovery rates (FDRs) determined via Benjamini-Hochberg correction of associated *P*-values. Gene ontology analysis was performed on uLIPSTIC-correlated (FDR < 0.05, Spearman r > 0) or anticorrelated genes (FDR < 0.05, Spearman r < 0) using goseq (v1.50.0) with ontology terms from Gene Ontology: Biological Process (GO:BP) and significance cutoff of FDR < 0.05^3^. To minimize redundant terms, GO:BP terms were ranked by proportion of overlap with up-or downregulated genes, then any term that itself had >70% gene overlap with a preceding term was removed.

### Fluorescence activated cell sorting and analysis

Cell suspensions were incubated with the appropriate antibodies diluted in FACS buffer for 30 minutes at 4°C and washed with FACS buffer. For staining of intracellular targets, cells were fixed with Fixation/Permeabilization solution (eBioscience™ Foxp3 / Transcription Factor Staining Buffer Set) for 30 minutes at 4°C. Cells were then incubated with antibodies for intracellular targets diluted in Permeabilization buffer (from the same kit as above) for 30min at 4°C. For EdU staining, the Click-iT™ EdU Cell Proliferation Kit for Imaging, Alexa Fluor™ 647 dye from ThermoFisher was used according to maufacturer’s instructions. To stain dead cells, samples were incubated with DAPI (0.2 μg/ml) right before acquisition/sorting. Alternatively, for fixed samples, cells were stained with the Fixable Viability Dye (ThermoFisher Scientific, 1:1000) for 15 minutes at 4°C prior to fixation. Below the antibodies and gating strategies for the main cell types analyzed in the study are outlined:

#### Epithelial skin stem cells

The following antibodies were used: CD49f (α6) BV510 (GoH3, BD Biosciences, 1:200); CD200 PE (OX-90, Biolegend, 1:200) or APC-R700 (OX-90, BD Biosciences, 1:100); Sca1 PerCP-Cy5.5 (D7, Biolegend, 1:200); CD34 FITC or BV421 (RAM34, BD Biosciences, 1:100), LTBR PE (5G11, Biolegend, 1:200), I-A/I-E (MHC-II) PE/Dazzle 594 (M5/a, Biolegend, 1:200), Galectin-3 PE-Cy7 (M3/38, Biolegend, 1:200), CD90.1 PE-Cy7 (OX-7, Biolegend, 1:200). Different stem cell population were defined as follows: IFE-SC (live, α6^high^, CD200^neg^, Sca1^+^); uHF-SC (live, α6^high^, CD200^+^, Sca1^neg^, CD34^neg^); bulge SC (live, α6^high^, CD200^+^, Sca1^neg^, CD34^+^)

#### Immune cells

The following antibodies were used: CD45 AlexaFluor700 (30-F11, Biolegend, 1:200), TCR beta chain PerCP-Cy5.5 (H57-597, Biolegend, 1:200), TCRγδ BV421 or Alexa488 (GL3, Biolegend, 1:200), CD90.2 Alexa488 or PE-Cy7 (30-H12, Biolegend, 1:200), TCR Vγ5 APC or PE (536, Biolegend, 1:200), LAG-3 Alexa647 or PE-Cy7 (C9B7W, Biolegend, 1:200), IL-17 PE or PE-Cy7 (TC11-18H10.1, Biolegend, 1:200), IFNγ APC (XMG1.2, Biolegend, 1:200), CD4 PE-Cy7 (GK1.5, 1:200 Invitrogen), CD8 PE (53-6.7, 1:200, Biolegend), CXCR3 APC (CXCR3-173, Biolegend, 1:200), CD11b PE-Cy7 (M1/70, 1:200, Biolegend). For AREG intracellular cytokine staining, cells were incubated first with the polyclonal goat anti-mouse AREG antibody (1:200 RnD Systems), followed by donkey anti-goat Al488 or Al647 secondary antibody (1:300, Life Technologies). For intracellular cytokine staining, cells were incubated for 3h at 37°C in RPMI+10%FBS with 1μg/ml Ionomycin (BRAND), 100μg/ml PMA (BRAND) and BD Golgi plug (1:100, BD Biosciences). Different immune populations were defined as follows: DETC (live, CD45^+^, CD90^+^, TCRβ^neg^, TCRγδ^high^ or Vγ5^+^ when applicable); TCRαβ+ T cells (live, CD45^+^, CD90^+^, TCRβ^+^, TCRγδ^neg^); ILCs (live, CD45^+^, CD90^+^, TCRβ^neg^, TCRγδ^neg^).

#### uLIPSTIC experiment

Biotinylated cells were detected using anti-biotin PE (Bio3-18E7, 1:50 Miltenyi Biotec). For single cells sequencing, a streptavidin-PE TotalSeq-C0961 barcoded antibody (Biolegend, 1:1000) was used.

Labelled cells were analysed using LSRFortessa Analyzer (BD Biosciences) and sorted with Sony Biotechnology MA800 or with BD FACSAriaII (equipped with Diva Software v.8.0) cell sorters. Analysis was carried out using with FlowJo program.

### Immunofluorescence image analysis

Whole mount confocal images were acquired using an inverted Dragonfly 202 spinning disk confocal system (Andor Technology Inc.) using a 20x air objective and a Zyla camera. Four laser lines (405, 488, 561 and 625) were used for near simultaneous excitation of DAPI, Alexa-488, RRX and Alexa-647 fluorophores. Tiled stacks of 1-2.5μm were acquired using the Andor Fusion Software (v2.3). Images were stitched and aligned using Imaris Stitcher (Bitplane) software using DAPI as a fiducial. Imaris software was used to perform image analysis and create representative figures using max projection of multiple stacks (Extended Sections option). Scale bars are indicated in each figure.

#### Quantification of LTB^+^ cells

to quantify LTB^+^ DETC distribution in the skin, extended sections (max projection of multiple stacks) of the IFE and the upper hair follicle compartment were visualized and analysed separately. All the LTB^+^ DETC (TCRγδ^+^ and LTB-tdT^+^) in each field of view (FOV) were counted manually and the percentage of LTB^+^ DETC in each compartment (i.e. IFE or uHF) out of the total number of LTB^+^ DETC were calculated and plotted. The area at the border between hair follicles and IFE was defined as “junction” and was removed from the analysis. For each mouse (represented by a dot in the plot), 3-9 FOVs were analysed.

#### Quantification of DETC numbers in anti-LAG3 treated mice

for quantification of total DETC abundance in the IFE, total number of DAPI^+^ and the total number of DETC (TCRγδ^+^) in a 3D FOV of the IFE compartment was quantified using the spot creation tool and spot rendering statistics tool (Imaris). DETC abundance in the IFE was then quantified as %DETC out of total number of DAPI^+^ cells in the IFE. Average number of total DAPI cells was comparable between the two experimental groups. For quantification of DETC in the uHF, total number of DETC in the uHF region and total number of hair follicles/FOV were counted manually using the extended section max projections tool. To calculate the average number of DETC/HF, the total number of DETC counted in each FOV was divided by the total of HF counted in the same FOV. For each mouse (indicated by a dot in the plot), 3-4 FOVs were quantified.

#### Quantification of CD8^+^ T cells/HF upon mosaic knockdown of LTBR

upper hair follicle region was visualized using max projection of multiple stacks (extended section option in Imaris software). Total number of CD8^+^ T cells/HF were manually counted and annotated. Hair follicles were then manually assigned to three categories based on their EGFP (marking lentivirus-transduced HF) intensity and distribution, being either completely negative, completely positive or only partially positive (i.e. a mix of transduced and untransduced cells in the same follicle). Partially positive follicles were excluded from the analysis. The average number of CD8^+^ cells/HF category was quantified. For each mouse, 4-5 FOVs were quantified.

### Keratinocytes proliferation

Keratinocytes isolated from postnatal pups were culture in E-low Ca media (Fuchs Laboratory^4,5^) in 96-well plates (black plastic, ThermoScientific). On day zero, keratinocytes were stimulated with different doses (25ng/ml, 50ng/ml, 250 ng/ml) of recombinant mouse Amphiregulin (Peprotech) or with vehicle (PBS). On day 4, media was removed and CellTiter-Glo (Promega) reagents were mixed 1:1 with media ad added to each well. Plates were covered with aluminium foil and incubated for 10 minutes at RT with horizontal agitation. Luminescence (proportional to the amount of ATP, peak emission 560nm) was acquired using Biotek Cytation 5 reader. Each condition was plated in technical triplicate.

### RNA extraction, cDNA reverse transcription and quantitative PCR

Cells to be processed for RNA extraction were lysed using RLT buffer (Qiagen). For intra-epithelial qPCR analyses, the skin intra-epithelial fraction (see above) was pelleted and resuspended in approximately 350μl of RLT buffer and immediately frozen and stored at −80°C for subsequent RNA extraction. For RNA extraction from specific cell types, cells were FACS-sorted directly into RLT buffer (maintaining a ratio 1:3 FACS sheath fluid:RLT buffer). An average of 200,000 cells/sample were used to isolate RNA. RNA was extracted using the Qiagen RNeasy Micro kit according to manufacturer’s instructions (including the DNase treatment step) and RNA was eluted using DNase/RNase-free water. Quality and concentration of RNA was assessed using Nanodrop instrument or with an Agilent 2100 Bioanalyzer (for bulk RNA sequencing).

For conversion to cDNA, the same amount of RNA (usually 100ng) was reverse transcribed using the Vilo Reaction mix/Superscript Enzyme mix (Invitrogen) according to manufacturer’s instructions.

To normalize cDNA amounts across samples, primers for *Ppib2* or *Hprt* were used as housekeeping genes. cDNA, specific primers (see below for the sequences) and SYBR green PCR MasterMix (Sigma Aldrich) were mixed and ran on a QuantStudio 6 Flex Applied Biosystems Fast Real-Time PCR system. Relative gene expression is plotted in the manuscript as log2 fold change compared to the average expression levels of control samples in each experiment (after normalization with housekeeping genes).

#### Primer sequences

*Ppib2:* GTGAGCGCTTCCCAGATGAGA; TGCCGGAGTCGACAATGATG

*Hprt:* TCAGTCAACGGGGGACATAAA; GGGGCTGTACTGCTTAACCAG

Areg: CAGTGCACCTTTGGAAACGA; ATGTCATTTCCGGTGTGGCT

Ki67: AGTCTCTTGGCACTCACAGC; ATTTTGTAGGGTCGGGCAGG

Nfkb2: CCTTCTCCTTGCCCCTGAAG; GTAGGCCAAGGAGGAGGAGA

Cxcl10: ATGACGGGCCAGTGAGAATG; AGCTAGGGAGGACAAGGAGG

Cxcl16: CAGTCCAAAAGCGTGTGTGG; CATGGCTGCAGTGAGGAAGA

Il33: TCGGCTGCTTGCTTTCCTTA; GTTCCCGGCCTCTTTGATCA

Ltb: ACCAGAAACTGACCTCAACCC; CTCAGAAACGCTTCTTCTTGGC

Ltbr: TGACATTGTGCTGGGCTTTG; TGCACACACTCATTGTCCAG

### LAG3 in vivo blockade and LTBR agonist administration

#### LAG-3 blocking antibodies

monoclonal antibodies against mouse LAG-3 (clone C9B7W, BioXcell) were injected intraperitoneally every other day (for a total of three injections) in wild-type C57BL/6 animals. For each treatment, 200μg in 150μl of PBS/mouse were injected.

As a control, mice were injected with the same volume of PBS. Two days after the last injection mice were injected with EdU and sacrificed 3 hours later for organ harvesting.

#### LTBR agonist treatment

wild type C57BL/6 mice ordered from JAX in second telogen were anaesthetized using isoflurane prior to intradermal injections with the LTBR agonist. Anaesthesia was maintained throughout the procedure using a nose cone for delivery of isoflurane. Half of the back skin was shaved using electric clippers and the surface sterilized using ethanol wipes. One millilitre insulin syringe with the needle bent to approx. 45° was used to shallowly inject the solutions through the epidermis and into the dermal space (at 3-5mm from the injection site). An injection volume of 25ul was delivered per injection site, for a total of two injections/mouse (one in the anterior and one in the posterior back). Injections were repeated every other day for a total of three times and mice were sacrificed one day following the last injection. Following each injection, mice were placed in their home cage on a heating pad to recover. To promote LTBR activation, an agonist monoclonal antibody targeting mouse LTBR (clone 3C8, Adipogen) was used. Each mouse received 7μg of antibody diluted in 50μl of PBS (25μl per injection site, 2 injection sites in total). Treatment was repeated for three times in total (each time with 7μg of antibody). Control mice were injected with anti-mouse IgG1 isotype control (at the same dose). Mice were randomly assigned to the two experimental groups.

### Osmotic pump for rAREG systemic delivery

For systemic delivery of recombinant mouse AREG (Peprotech), Alzet micro-osmotic pumps (Model 1003D) were used. Second telogen wild type female mice were anesthetized and their back sterilized for surgery. A small incision on the right side of the mouse back was created with scissors and the pumps were inserted in the subcutaneous space. The incision was then closed with surgical staples. For rAREG experiments, a three-day long delivery pump was used, loaded with 35 μg of rAREG or PBS as a control. The contralateral (i.e. left) side of the mouse back was used for analysis four days after the surgery.

### Bulk RNA sequencing

In the LTBR-agonist treatment experiment, uHF-SC were sorted directly in RLT buffer (Qiagen) and RNA was extracted following the protocol described above. All samples had RNA integrity (RIN) numbers >8.2 (as measured with an Agilent 2100 Bioanalyzer).

Comparable amounts of RNA per sample were used to prepare bulk RNA-sequencing libraries using NEB Ultra II Directional RNA Library prep with dual indexing (plus Poly A isolation module) following manufacturer’s guidelines. Libraries were then uniquely barcoded, pooled and sequenced on an Illumina Novaseq 6000 – S4 Flow cell using paired end reads (at Weill Cornell Medical College’s Genomic Core Facility).

RNA-seq transcript quantification from raw 40bp paired-end sequencing reads was performed with salmon (v1.10.3), using selective alignment of reads via the --validateMappings parameter^6^. Raw counts were used as input into DESeq2 (v1.42.1)^7^ for differential gene expression testing with default parameters. Gene ontology analysis was performed on upregulated (DESeq2 *P-adj* < 0.05, log2FoldChange > 0) or downregulated genes (DESeq2 *P-adj* < 0.05, log2FoldChange < 0) in upper hair follicle stem cells exposed to LTBR agonist vs. isotype control using goseq (v1.50.0) with ontology terms from Gene Ontology: Biological Process (GO:BP) and significance cutoff of FDR < 10^-4 3^. To minimize redundant terms, GO:BP terms were ranked by proportion of overlap with up-or downregulated genes, then any term that itself had >70% gene overlap with a preceding term was removed.

### Transwell migration assay

#### Stem cells supernatant preparation

Skin epithelial stem cells from different compartments (i.e. IFE, uHF and Bulge) were FACS-sorted as described above and maintained in culture on a layer of mitomycin C inactivated 3T3/J2 feeder fibroblast cells (gift from Howard Green, now available at Kerafast) with SY media (Fuchs Lab, previously described in^8^, Y-27632 from Sellechem). At day −3 and −1, stem cells were stimulated with 1μg/ml of LTBR-agonist (3C8, Adipogen) or Isotype control (anti-IgG1, Adipogen). At day 0 (i.e. when the transwell migration assay was performed), the supernatant from the stem cell cultures was collected into 1.5ml Eppendorf tubes and spun down at 1800rpm for 5 minutes. The cell-free supernatant was then collected and transferred to another tube, before being plated in the lower transwell chamber.

#### Activated lymphocytes preparation

at day −3, one spleen was harvested from a wild type C57BL/6 mouse and the red blood cells were lysed using RBC lysis buffer 10x (Biolegend) diluted in water for 5 minutes on ice. Cells were then washed with FACS buffer and spun down at 1500rpm for 5 minutes. Splenocytes were then resuspended in 14-20 ml of RPMI+10% FBS+ 1% penicillin and streptomycin (P/S). Cells were then stimulated in a 96-well U-bottom plate, each well containing 200μl of splenocytes + 1.5-2μl of Dynabeads Mouse T-activator CD3/CD28 (Gibco). Approximately 30-50 wells were plated and cells were incubated at 37°C with 7.5% CO_2_. On day 0, splenocytes in all the wells were pooled together, magnetic beads were removed using a magnet and cells were spun down at 1500rpm for 5min and resuspended in fresh RPMI+10% FBS+1%P/S media.

#### Transwell assay

the HTS Transwell 96 permeable support with 5.0μm pore (Corning) was used for the assay. In the lower chamber, 170μl of supernatant from cultured stem cells (collected as described above) were plated. When indicated, 50μg/ml of InVivoMab anti-mouse CXCL10 (IP-10, BioXcell) was added to the lower chamber. After all the lower chambers were plated, the transwell insert was adjusted on top of the plate and 100μl of splenocytes (equivalent to approx. 500.000 cells, prepared as described above) were plated in the upper chamber. Cells were allowed to migrate for 90 minutes at 37°C with 7.5% CO_2_. At the end of the incubation, the transwell insert was removed and the splenocytes migrated to the lower chamber were collected and stained with fluorophore-conjugated anti-CD45, CD4, CD8 and CXCR3 antibodies for subsequent FACS analysis. Ten microlitre of counting beads (Sperotech Inc., equivalent to 10.000 beads), were added to each sample right before acquisition at the LSRFortessa analyzer.

### In utero lentiviral injection and Cas9-mediated LTBR deletion

To identify efficient CRISPR single guide RNAs (sgRNA) against mouse LTBR, we synthesized oligonucleotides targeting exon 3 or exon 10 with BsmBI restriction sites at 5’ and 3’ respectively (sequences reported below). Oligonucleotides were subcloned into a pLKO-Cre stuffer v4 (Addgene, Daniel Schramek lab^9^), following the Schramek lab protocol. High-titre lentivirus was prepared and E9.5 embryos were infected with lentiviruses delivered by ultrasound-guided microinjection into the amniotic sac as previously described^10^. To select a guide for functional experiments, efficient silencing in uHF-SC upon E9.5 lentiviral delivery was assessed in adult mice by flow cytometry staining of EGFP and LTBR.

#### Single guide RNA sequences

1. FW: CACCGTTCATTATAGGAATTATGG; REV: AAACCCATAATTCCTATAATGAAC
2. FW: CACCGCTGTCTACACCCTACCAGG; REV: AAACCCTGGTAGGGTGTAGACAGC

### S.epidermidis colonization

*Staphylococcus epidermidis* (strain NIHLM087, obtained from J. Segre) was grown in tryptic soy broth (TSB, Millipore) overnight at 37 °C and single bacterial colonies were grown the day after on a Heimplate Mannitol Salt Agar (MSA, Millipore) plate. Every other day, one bacterial colony was grown overnight in 30ml of TSB at 37 °C. The following day, bacterial cultures were spun down at 4300rpm for 10minutes and resuspended in 30ml of fresh TSB for 5-7 hours at 37 °C. Bacterial were spun down again and resuspended in sterile PBS. Each mouse was shaved with electric clippers and associated with 1ml of bacterial suspension (∼10^9^ colony-forming unit [CFU]/mL) across the entire skin surface of the mouse back using a sterile cotton swab. Applications were repeated every other day for a total of four applications. One-two weeks after the last application, the back skin was swabbed with a sterile cotton swab pre-wetted in sterile PBS. MSA plates were streaked with the swabs and bacterial colonies were grown overnight at 37 °C. One day later, colonies on each plate (representing each mouse) were counted. When colonizing Ltb-CreER x R26-DTA and R26-DTA control mice, littermate controls were used. Five-seven weeks old mice mice were separated in different cages based on genotype, treated with tamoxifen over the course of two weeks and one-two weeks after the last tamoxifen treatment mice were colonized with *S.epidermidis*.

### AREG ELISA on intra-epithelial tissue

A tissue biopsy of approximately 1.5 cm^2^ was collected from the mouse back skin and the dermal tissue was removed by scraping with a dull sclapel. The tissue was placed in an eppendorf, snap frozen in liquid nitrogen and stored at −80°C until later use. To isolate the protein extract, frozen tissues were crashed using a metal hammer and a metal block. The tissues pieces were collected in an eppendorf and lysed by bead beating (0.9-2mm diameter stainless steel beads, Next Advance) using a lysis buffer containing 1x RIPA buffer (Millipore) + 0.1% SDS + 1:100 HALT protease inhibitor (Thermo Fisher) + 100μM phenyl methyl sulfonyl fluoride (Thermo Scientific) + 1 mM benzamidine hydrochloride (Sigma). Total protein concentration was quantified using a BCA protein assay (ThermoScientific) and all the samples were diluted to the same concentration (1mg/ml) of total protein. AREG concentration was quantified by ELISA (Mouse AREG ELISA Kit from Thermo Fisher Scientific) following manufacturer’s instructions.

### Statistics and reproducibility

All data from each experiment were included in the analyses unless positive or negative controls failed to work, in which case the experiment was removed from analysis. Mice were assigned randomly to experimental groups and no statistical methods were used to predetermine sample size. Given the unanbiguity in phenotypes and readouts or and in the use of internal control, experiments were not performed blindly. Statistical analyses were performed using GraphPad Prism (version 10.6.1). Sample size, statistical tests and replicates are indicated oin each figure legend. Unless stated otherwise, experiments comparing two experimental groups were analyzed using (paired or unpaired) two-tailed Student’s t-test with a 95% confidence interval. When comparing three or more groups, oridinary one-way ANOVA with Tukey’s multiple comparisons test. Data are visualized as scatter plots with or without bars or as individual values connected with lines for paired analyses.

